# The kinetics and mobility of a ParA ATPase drive carboxysome distribution in *Halothiobacillus neapolitanus*

**DOI:** 10.64898/2026.03.03.709431

**Authors:** Christopher A. Azaldegui, Hema M. Swasthi, Longhua Hu, Lisa T. Pulianmackal, Hannah Rivett-Trznadel, Jian Liu, Anthony G. Vecchiarelli, Julie S. Biteen

## Abstract

Carboxysomes are bacterial microcompartments that drive efficient carbon fixation in autotrophic bacteria. Critical to their function and inheritance is their spatial organization by the ParA-type ATPase, McdA, and its partner protein, McdB. Here, we investigate the α-carboxysome McdAB system in *Halothiobacillus neapolitanus* using biochemical assays, quantitative fluorescence imaging, and mathematical modeling. We find that, unlike most ParA-type ATPases, the ATPase activity of McdA is only stimulated by DNA rather than by its partner protein McdB. Despite this difference, McdB conserves the ability to displace McdA from DNA, suggesting that ATP hydrolysis and DNA unbinding by McdA are not strictly coupled. Together with its ability to diffuse while bound to DNA, McdA forms gradients on the nucleoid that prevent carboxysome aggregation via a Brownian ratchet mechanism. Overall, these findings reveal key differences in a ParA-type ATPase that may be specific for the spatial organization of protein-based organelles in bacteria.

## INTRODUCTION

Efficient cellular metabolism is achieved by compartmentalizing enzymatic reactions. Prokaryotes achieve this process by using bacterial microcompartments (BMCs), ubiquitous organelles that encapsulate metabolic pathways within a permeable protein-based shell (Shively et al., 1973; Kerfeld et al., 2018; Sutter et al., 2021). The most extensively studied BMC is the carboxysome, which selectively concentrates the enzyme ribulose-1,5-biphosphate carboxylase/oxygenase (RuBisCO) with CO_2_ to efficiently catalyze carbon fixation in many autotrophic bacteria (Shively et al., 1973; Yeates et al., 2008). Carboxysomes exist as either α or β types, differing in structural composition, assembly, and regulation (Yeates et al., 2008; Rae et al., 2013; Kerfeld and Melnicki, 2016; Liu, 2022). Both types are found in cyanobacteria, while α-carboxysomes are also widespread across other bacterial phyla and more closely resemble other types of BMCs (Kerfeld and Melnicki, 2016; Sutter et al., 2021). Therefore, understanding α-carboxysome regulation is crucial for comprehending broader BMC biology.

An important aspect of carboxysome regulation is their spatial organization within the cell. In *Synechococcus elongatus* PCC 7942, β-carboxysomes are positioned along the nucleoid by the Maintenance of Carboxysome Distribution (Mcd) system, consisting of McdA and McdB (Savage et al., 2010; MacCready et al., 2018; MacCready and Vecchiarelli, 2021). McdA is a DNA-binding deviant Walker-like ATPase similar to the ParA/MinD ATPase family that bacteria employ to organize genetic- and protein-based cellular complexes (Jalal and Le, 2020; Salomon et al., 2021; Pulianmackal and Vecchiarelli, 2024). McdA binds DNA non-specifically and forms gradients on the nucleoid in the presence of its partner protein McdB (MacCready et al, 2018). McdB is an adaptor protein that localizes to carboxysomes and stimulates the ATPase activity of McdA, resulting in the release of McdA from DNA (MacCready et al., 2018; Basalla et al., 2023; Basalla et al., 2024). This repeated cycle of interactions between DNA, McdA, and McdB results in the equal distribution of carboxysomes, ensuring their even inheritance after cell division (Savage et al., 2010; MacCready et al., 2018; Rillema et al., 2021).

Among the best understood ParA/MinD ATPase family proteins are the ParABS systems involved in plasmid and chromosome partitioning (Jalal and Le, 2020; Azaldegui et al., 2025). ParA binds DNA non-specifically in its ATP-bound dimer state, and its partner protein, ParB, binds to a specific DNA sequence (*parS*) to form a nucleoprotein complex. ParB stimulates the release of ParA from the nucleoid, creating gradients that drive the directed and persistent motion of DNA cargos towards increased concentrations of ParA. This interplay of interactions results in evenly spaced positioning of cargos within cells (e.g., one copy at the mid-cell, two copies at the quarter positions).

Given its similarities to ParABS systems, we proposed that McdAB distributes β-carboxysomes via a Brownian ratchet mechanism (MacCready et al., 2018; Hu et al., 2017; Jalal and Le, 2020; Azaldegui et al., 2025). In our Brownian ratchet model (Hu et al., 2015; Hu et al., 2017; Hu et al., 2021), ParB-stimulated release of ParA from DNA and the subsequent stalled relocalization of ParA to these regions result in a ParA-depletion zone around the cargo. The asymmetry of this ParA distribution on the nucleoid drives directional movement of the cargo through the remaining ParA-ParB bonds. In the presence of multiple cargo copies, this process results in their directed segregation and subsequent equidistant positioning on the nucleoid. While several other Brownian-ratchet models have been proposed to explain ParA-mediated cargo positioning, they are all fundamentally predicated on the same biochemical cycle of ParA ATPase activity and the resulting asymmetric distribution of ParA dimers surrounding the cargo (Vecchiarelli et al., 2013; Vecchiarelli et al., 2014; Lim et al., 2014; Surovtsev et al., 2016; Le Gall et al., 2016; Walter et al., 2017; Kohler et al., 2022).

Recently, we identified an α-McdAB system in *Halothiobacillus neapolitanus*, which positions α-carboxysomes and is widely encoded in α-carboxysome-containing proteobacteria (MacCready and Tran et al., 2021). Unlike β-McdA, α-McdA preserves the full deviant Walker A box and is predicted to be structurally homologous with typical ParA ATPases (MacCready and Tran et al., 2021; Pulianmackal et al., 2023). The existence of α-McdAB-like sequences within and near operons of other types of BMCs suggests that *H. neapolitanus* α-McdAB is a model for BMC spatial regulation in general (MacCready and Tran et al., 2021). While the functions of *H. neapolitanus* α-McdAB are known, the biochemical properties of α-McdA remain unclear. Furthermore, measuring the biochemical and biophysical properties of the α-McdAB system, which are critical for our Brownian ratchet model, will clarify the extent to which this model can explain carboxysome and BMC positioning in general.

Here, we combined biochemical assays, quantitative fluorescence microscopy, and single-molecule imaging to determine whether the biochemical and biophysical properties of the McdAB system in *H. neapolitanus* are consistent with the Brownian ratchet model. We demonstrate that gradients of McdA on the nucleoid result in the confined motion of distributed carboxysomes. Additionally, we discovered that α-McdA ATPase activity is significantly stimulated by DNA alone, and this activity is not further stimulated by its partner protein, α-McdB. However, in agreement with our Brownian ratchet model, α-McdB still promotes α-McdA release from DNA, suggesting that ATP hydrolysis and DNA release are not strictly coupled. Furthermore, we demonstrate that McdA can diffuse while bound to the nucleoid, possibly through transient DNA interactions, providing a mechanism for McdA relocalization independent of McdB. Our experiments, along with Brownian ratchet simulations based on biochemically and physically determined parameters, indicate that McdAB functions as an anti-aggregation system that maintains the separation of carboxysomes rather than positioning them at specific locations within the cell.

## RESULTS

### Carboxysomes are evenly distributed in H. neapolitanus

To quantify the α-carboxysome (carboxysome hereafter) distribution in *H. neapolitanus*, we imaged fluorescently labeled carboxysomes (CbbS-mTurquoise2) (Figure 1a). We calculated carboxysome density as the number of carboxysome foci per cell length (Figure 1b) and measured 2.7 ± 0.6 foci/µm length of the cell (mean ± standard deviation, SD) (Figure 1b). In contrast, Δ*mcdA* cells have a reduced carboxysome density of 0.8 ± 0.2 foci/µm (Figure 1b), indicating aggregation of carboxysomes at the cell poles (Figure 1a). We also quantified carboxysome distribution by measuring the spacing between each carboxysome focus and its nearest neighbor focus (Supplementary Fig. 1). In WT cells, carboxysome foci are on average 0.31 ± 0.14 µm apart, while in Δ*mcdA* cells with more than one focus, these are spaced by 1.15 ± 0.4 µm, which indicates nucleoid-excluded foci at both cell poles (MacCready and Tran, et al., 2021). Notably, carboxysome foci are staggered in WT cells (Figure 1a; white arrows); a feature also observed in *S. elongatus* cells with higher carboxysome numbers (MacCready et al., 2018), which may be due to the McdAB system maximizing inter-carboxysome spacing.

**Figure 1:**
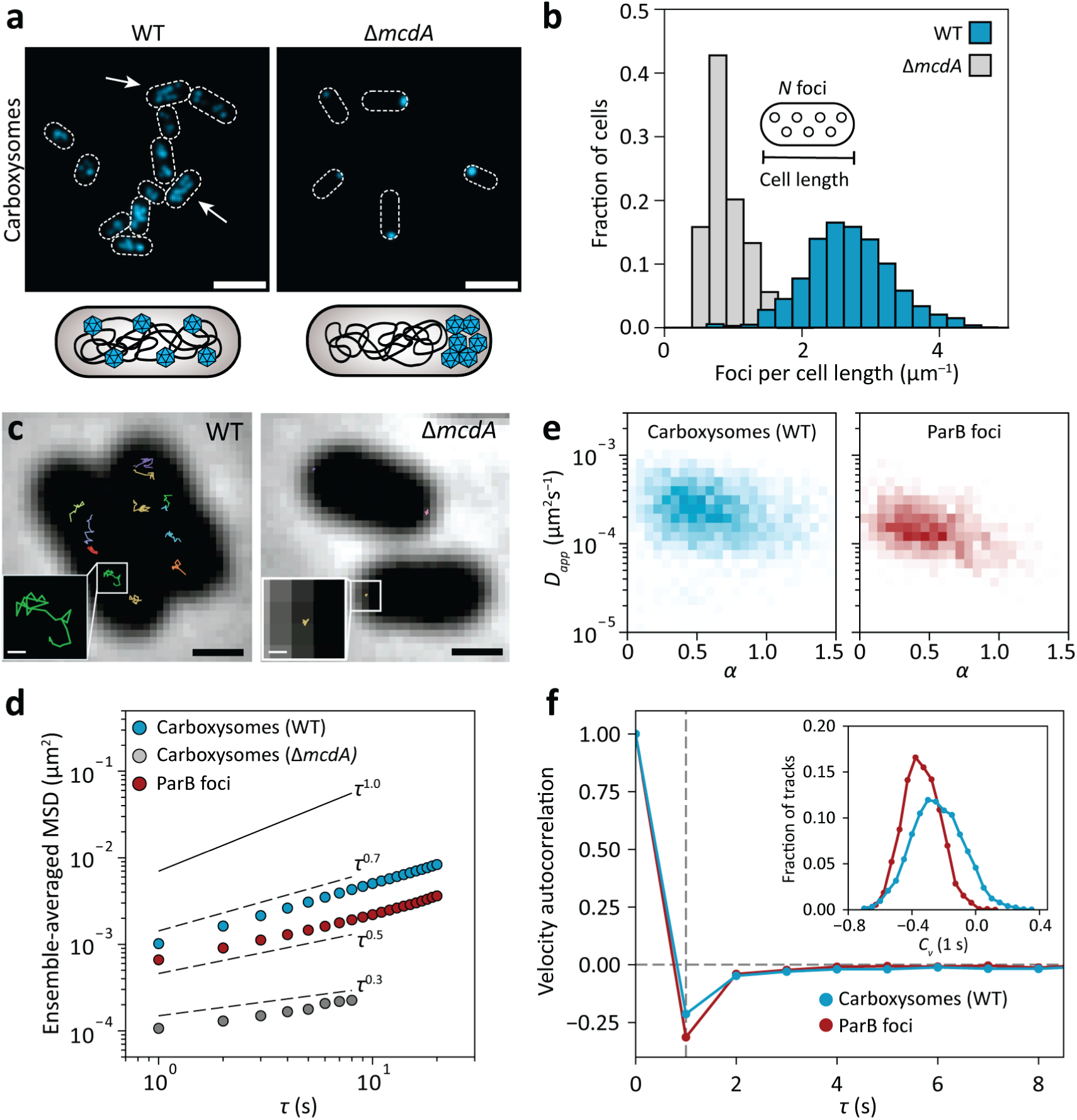
Carboxysomes in *H. neapolitanus* are dynamic and distributed. **a.** Representative fluorescence images of wildtype (WT, left) and Δ*mcdA* (right) *H. neapolitanus* cells (white dashed outlines) expressing the carboxysome marker, CbbS-mTurquoise2 (cyan). White arrows: cells with staggered carboxysomes. Scale bars: 2 µm. **b.** Distributions of linear carboxysome focus density in WT (*N* = 1061 cells) and Δ*mcdA* (*N* = 1106 cells) from two independent datasets. **c.** Representative trajectories of single carboxysome particles in WT (left) and Δ*mcdA* (right) cells. Scale bars: 500 nm. Insets: Zoom-in of a carboxysome trajectory. Scale bars: 50 nm. **d.** Population average mean-squared displacement (MSD) vs. lag time, *τ*, curves for carboxysome trajectories in WT (*N* = 3630 trajectories) and Δ*mcdA* (*N* = 1896 trajectories) cells and ParB-mNG chromosome foci (*N* = 1284 trajectories), from at least two independent datasets per sample. Solid lines: Brownian motion (*α* = 1). Dashed lines: the average *α* for each sample. **e.** 2D histograms of the *D_app_* and *α* exponent for the trajectories of carboxysomes (*N* = 3630 trajectories) and ParB-mNG chromosome foci (*N* = 1284 trajectories) in WT cells. **f.** Average velocity autocorrelation function for trajectories of carboxysomes (*N* = 3680 trajectories) and ParB-mNG foci (*N* = 1304 trajectories) in WT cells as a function of *τ*. The vertical dashed line at *τ* 1 s demarcates the sampling time of the experiment. The inset shows the distribution of velocity autocorrelation values (*C_v_*) at 1 s

### Carboxysomes exhibit confined motion on the nucleoid

The tight distribution of inter-carboxysome spacing led us to investigate the diffusive dynamics of carboxysomes to determine whether they exhibit directed motion or local excursions on the nucleoid. Using time-lapse fluorescence microscopy at 1-second intervals, we tracked carboxysome motion inside live *H. neapolitanus* cells (Figure 1c). In Δ*mcdA* cells, carboxysomes form static aggregates at cell poles, serving as an immobile control (Figure 1c). We fit the mean-squared displacement (MSD) of each track across various time lags (*τ*) to the diffusion model MSD = 4*D_app_ τ^α^*, where *D_app_* is the apparent diffusion coefficient of the particle and *α* is the anomalous diffusion exponent. For WT cells, *D_app_*is (2.4 ± 0.05) × 10^−4^ µm^2^s^−1^, over an order of magnitude higher than the immobile carboxysome foci in Δ*mcdA* cells, (0.07 ± 0.01) × 10^−4^ (Figure 1d and Supplementary Fig. 2). Notably, 27% of carboxysomes in Δ*mcdA* cells diffuse faster with *D_app_* = (0.23 ± 0.02) × 10^−4^ µm^2^s^−1^, which we interpret as smaller carboxysome clusters or individual carboxysomes diffusing, distinct from the large immobile cluster of carboxysomes at the cell pole (MacCready and Tran et al., 2021).

Furthermore, carboxysomes generally exhibit an anomalous diffusion exponent, *α*, less than 1, indicating subdiffusive motion (Figure 1e). We hypothesized that the carboxysomes are tethered to DNA via McdA-McdB interactions (MacCready et al., 2018; MacCready and Tran et al., 2021). To test if *α* correlates with chromosome behavior (Weber et al., 2010; Javer et al., 2013; Lim et al., 2014), we tracked the chromosomal partition complex by imaging ParB-mNeonGreen (ParB-mNG) in *H. neapolitanus* under the same conditions. ParB-mNG foci showed subdiffusive behavior (Figure 1e), with a tight *α* distribution centered approximately at 0.4, consistent with chromosomal locus behavior in *Escherichia coli* and *Vibrio cholerae* (Weber et al., 2010; Javer et al., 2013). A small subset of the ParB-mNG foci has a larger *α*, which may indicate freer chromosomal movements (Javer et al., 2014) or segregating chromosomes, as many of the cells imaged had two ParB-mNG foci. Both ParB-mNG foci and carboxysomes have a negative velocity autocorrelation at a 1-second timescale (Figure 1f), reflecting the viscoelastic cytoplasm (Parry et al., 2014). The absence of directional motion, oscillatory behavior, or random diffusion of carboxysomes and the ParB partition complex suggests that both cargos behave elastically, consistent with other reports of ParA-mediated positioning (Ietswaart et al., 2014; Lim et al., 2024; Köhler et al., 2022). However, the faster mobility and less constrained motion of carboxysomes (Figure 1d-f) relative to the chromosome suggest additional driving factors. For instance, the motion of β-carboxysomes in *S. elongatus* is confined, but they move towards regions of increased McdA concentrations on the nucleoid (Savage et al., 2010; MacCready et al., 2018; Sun et al., 2019; Dudley et al., 2025), indicative of a chemophoretic force (Vecchiarelli et al., 2013; Hu et al., 2015; Hu et al., 2017; Hu et al., 2021). Other possibilities include intrinsic carboxysome diffusion while untethered from the nucleoid (Hu et al., 2021) and DNA-based cargo relaying (Lim et al., 2014; Surovtsev et al., 2016b).

### Construction and characterization of functional McdA fluorescent fusions

To investigate the driving factor for carboxysome dynamics, we sought to visualize functional α-McdA (McdA hereafter) *in vivo*. We constructed a PAmCherry-McdA (PAmC-McdA) fusion with a GSGSGS linker, replacing the native *mcdA* gene. Carboxysome localization served as the phenotypic indicator of McdA function (Figure 1a and Supplementary Fig. 3a-b). Unlike the *S. elongatus* homolog (MacCready et al., 2018), this N-terminal fusion causes carboxysomes to mislocalize, forming a single carboxysome focus that is randomly positioned along the cell length, suggesting PAmC-McdA can load carboxysomes onto the nucleoid but not distribute them (Supplementary Fig. 3c). We also generated a C-terminal McdA-PAmC fusion, commonly used in functional ParA fluorescent fusions (Schofield et al., 2010; Ptacin et al., 2010; Lim et al., 2014), but this fusion causes the carboxysomes to aggregate at the cell poles (Supplementary Fig. 3d).

In some instances, fluorescent proteins have been fused internally to target bacterial proteins without perturbing function (Bendezu et al., 2008; Moore et al., 2016; Bisson-Filho et al., 2017; Cameron and Margolin, 2023). We examined the predicted AlphaFold2 (AF2) structure of McdA (Pulianmackal et al., 2023) and identified a putative disordered region with low prediction confidence between residues 55 and 64 (Supplementary Fig. 3e). Only residue Q62 within this loop is predicted to be functionally significant, and the loop is not conserved among other ParA/MinD models, suggesting that it is not critical for ParA/MinD function. We inserted PAmCherry and mNeonGreen between McdA residues 59 and 60, flanking them with GSGSGS linkers to create the sandwich (SW) fusions *mcdA-PAmC^SW^* and *mcdA-mNG^SW^* (Supplementary Fig. 3f). Complementation of these fusions in Δ*mcdA* cells restored carboxysome distribution (Supplementary Fig. 3g-h), with only a slightly reduced carboxysome focus density: we measured approximately 2.2 ± 0.6 and 2.4 ± 0.6 carboxysome foci per micron cell length for McdA-PAmC^SW^ and McdA-mNG^SW^, respectively, compared to the 2.7 ± 0.6 measured in unlabeled McdA cells (Figure 1b and Supplementary Fig. 3i). We also measured comparable mobility between carboxysomes across all functional strains, further supporting the functionality of the McdA sandwich fusions (Supplementary Fig. 3j).

### McdA forms gradients on the nucleoid independent of its McdB partner

We studied the spatial distribution and dynamics of McdA by time-lapse fluorescence imaging. McdA-mNG^SW^ did not exhibit the pole-to-pole oscillations observed for McdA in *S. elongatus* (Figure 2a-b). Instead, McdA-mNG^SW^ proteins form high-density regions on the nucleoid, resulting in short-scale gradients (Figure 2a-b and Supplementary Fig. 4a). This behavior is similar to that of the *E. coli* F plasmid ParA (Ah-Seng et al., 2013; Le Gall et al., 2016; Köhler et al., 2024). To determine whether these high-density regions are due to McdA binding to DNA, we imaged McdA[K20A]-mNG^SW^, which contains a mutation that inhibits DNA binding in other ParA-like proteins through ATP binding inhibition (Leonard et al., 2005). This mutant McdA is homogenously distributed, consistent with cytoplasmic localization and a lack of DNA binding (Figure 2a-b and Supplementary Fig. 4b).

**Figure 2:**
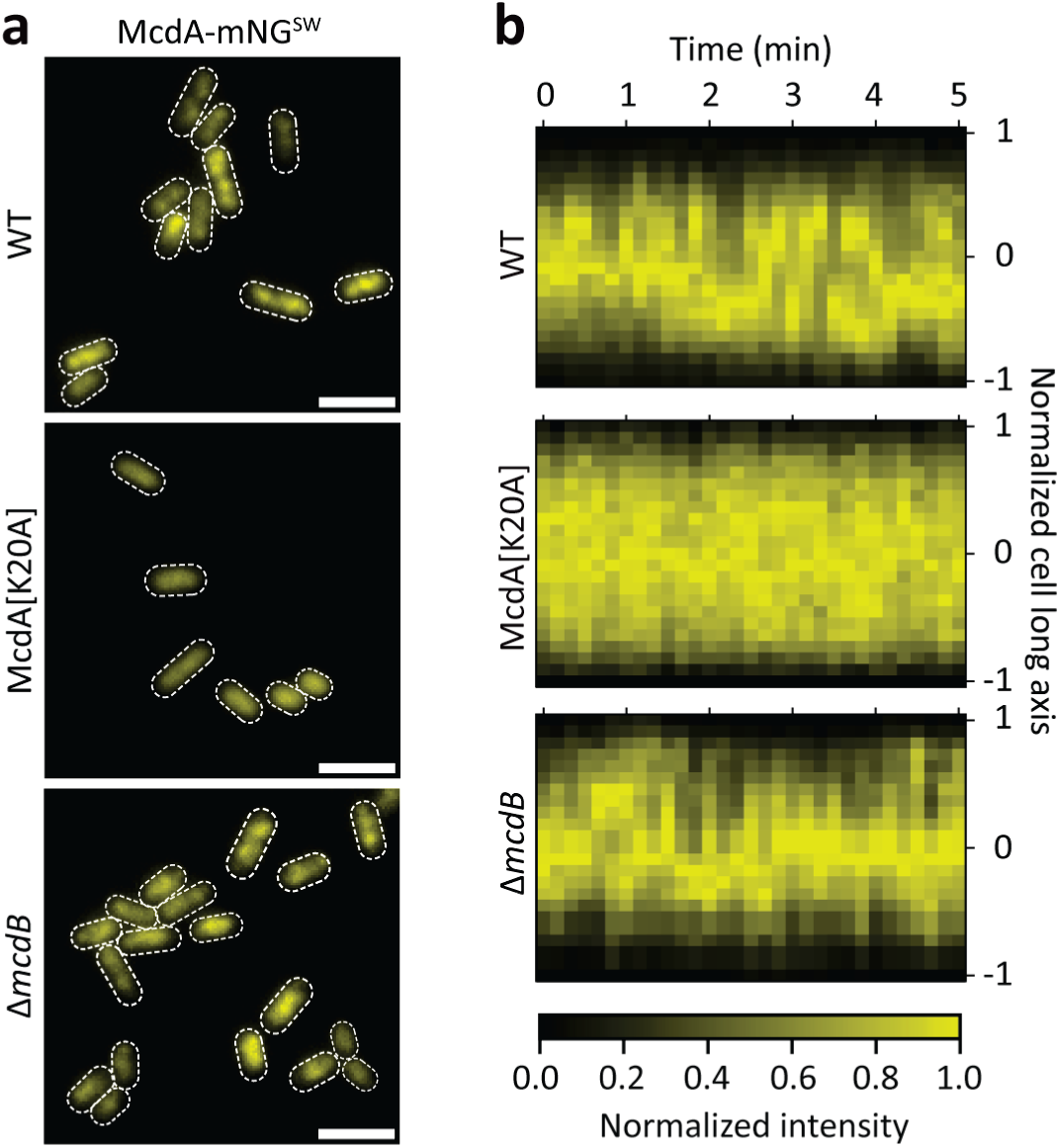
McdA forms gradients on the nucleoid. **a.** Representative images of McdA-mNG^SW^ in WT cells, McdA[K20A]-mNG^SW^ in WT cells, and McdA-mNG^SW^ in Δ*mcdB* cells. Dashed white lines are the cell outlines. Scale bars: 2 µm. **b.** Representative normalized kymographs across the cell long axis for McdA-mNG^SW^ in WT cells, McdA[K20A]-mNG^SW^ in WT cells, and McdA-mNG^SW^ in Δ*mcdB* cells.

While DNA binding by McdA enables the formation of these gradients, depletion of McdA on the nucleoid by McdA-McdB interactions (MacCready et al., 2018; Vecchiarelli et al., 2010) and stimulation of ATP hydrolysis may also influence the distribution of McdA, as is the case for a majority of the ParA/MinD partner proteins (Pulianmackal and Vecchiarelli, 2024). However, we imaged similar dynamics for McdA-mNG^SW^ in Δ*mcdB* cells as in WT cells (Figure 2a-b and Supplementary Fig. 4c). This unexpected result indicates that McdA can form gradients on the nucleoid independent of its association with McdB, suggesting differences in the biochemical properties of McdA relative to typical ParA ATPases.

### McdA binding to DNA is ATP- and dimerization-dependent

Building on the *in vivo* observations of McdA, we investigated the biochemical properties of McdA *in vitro*. We purified recombinant *H. neapolitanus* McdA and McdB proteins. We fused a SUMO tag (Kuo et al., 2014) to the N-terminus of McdA (*his-SUMO-mcdA*), which significantly increases its solubility (Supplementary Fig. 5). Without ATP, His-SUMO-McdA elutes as a single species in size-exclusion chromatography (Figure 3a), consistent with an McdA monomer. With ATP, the His-SUMO-McdA elution peak shifts and broadens (Figure 3a), indicating the presence of two species, likely the monomer and the ATP-dependent dimer form. The His-SUMO-α-McdA[K20A] mutant elutes as a single narrow peak both with and without ATP, matching the WT peak in the absence of ATP (Figure 3a). These results show McdA can dimerize upon ATP binding *in vitro*.

**Figure 3:**
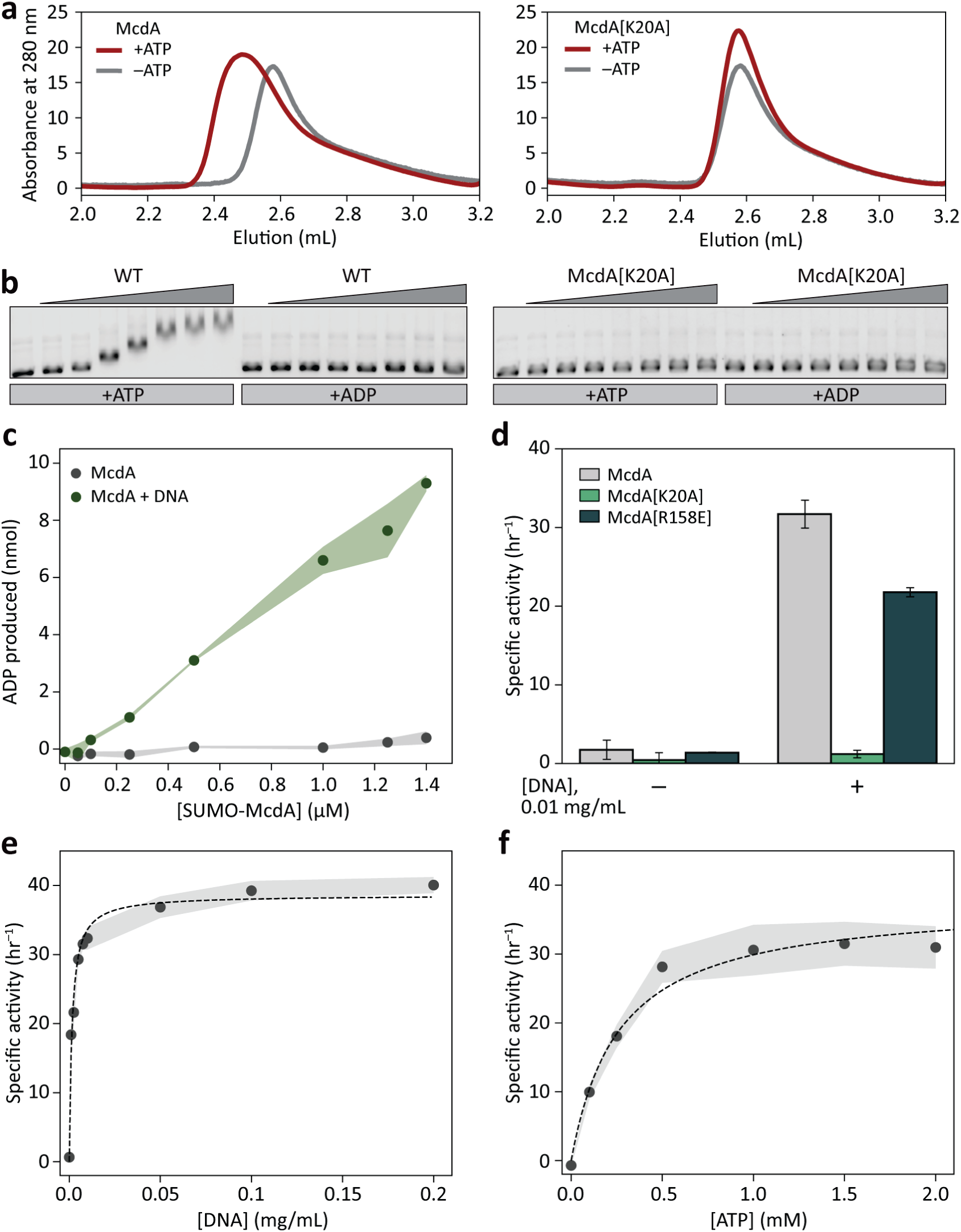
McdA binds DNA in its ATP-bound dimer state, and its ATPase activity is triggered by DNA. **a.** Gel filtration analysis of purified His-SUMO-McdA (left) and His-SUMO-McdA[K20A] (right) in the presence and absence of 1 mM ATP. **b.** Electrophoretic mobility shift assay (EMSA) with increasing concentrations (0, 0.5, 1, 2.5, 5, 10, 12.5, 14 µM) of His-SUMO-McdA (left) and His-SUMO-McdA[K20A] (right). 1 mM of ATP or ADP was used. **c.** ATPase activity of His-SUMO-McdA in the presence and absence of DNA (0.1 mg/mL). ATP concentration was kept fixed at 1 mM. **c.** Amount of ADP produced in reaction with 0.01 mg/mL DNA after 2 hr. Shading indicates the SEM of three independent experiments. **d.** Effect of DNA on ATPase activities of McdA, McdA[K20A], and McdA[R158E]. Error bars are the error on the linear regression of the mean ADP production rate (nmol hr^−1^) vs. the amount of McdA (nmol) from three independent experiments. **e.** Specific activity of His-SUMO-McdA (concentration fixed at 1.0 µM) measured as a function of DNA concentration (0 – 0.2 mg/mL). Shading indicates the SEM of three independent experiments. A Michaelis-Menten equation fit to the data (black dashed line) gives equilibrium constant, *K_DNA_*= 1.5 µg/mL and maximum rate, *V_max_* = 38.6 hr^−1^. **f.** Specific activity of His-SUMO-McdA (concentration fixed at 1.0 µM) as a function of ATP (0-2 mM) with fixed DNA concentration (0.01 mg/mL). Michaelis-Menten equation fit to the data (black dashed line) gives *K_M_* = 262 µM and *V_max_* = 37.7 hr^−1^.

Next, we probed the DNA-binding activity of McdA by Electrophoretic Mobility Shift Assay (EMSA). Adding His-SUMO-McdA to non-specific DNA (nsDNA) in the presence of ATP but not ADP causes an EMSA shift that correlates with His-SUMO-McdA concentration, demonstrating ATP-dependent DNA binding (Figure 3b). His-SUMO-McdA[K20A] does not shift the DNA mobility (Figure 3b), indicating that the non-specific DNA-binding activity of McdA depends on ATP binding and subsequent dimerization.

### McdA ATPase activity is triggered by DNA

*S. elongatus* β-McdA has strong ATPase activity compared to other ParA ATPases. We posited that this activity is due to its unique Walker A motif (SGGxxGKT), which differs from α-McdA and typical ParA ATPases (KGGxxGKT) (MacCready et al., 2018; MacCready and Tran et al., 2021; Hakim et al., 2021). This difference prompted us to test whether α-McdA acts as a robust ATPase like β-McdA (MacCready et al., 2018) or a poor ATPase like typical ParA ATPases (Davis et al., 1992; Ah-Seng et al., 2009; Lim et al., 2014; Schumacher et al., 2017). Unlike *S. elongatus* β-McdA, His-SUMO-McdA, in the presence of ATP, has a low specific activity of 1.7 h^−1^ (Figure 3c-d). However, the addition of nsDNA significantly increases the His-SUMO-McdA specific activity to 31.7 h^−1^ (Figure 3c-d), with maximum activity of *k_cat_* = 38.6 h^−1^ and a half-saturating DNA concentration *K_DNA_* = 1.5 µg/mL (Figure 3e). The rate also depends on ATP concentration (*K_M_*= 262 µM; Figure 3f). His-SUMO-McdA[K20A] has negligible ATPase activity in the presence and absence of DNA, as it does not bind ATP (Figure 3d). The large fold-change in ATPase activity of McdA in the presence of DNA is unusual for ParA/MinD ATPases, which generally have less than two-fold enhancements (Davis et al., 1992; Ah-Seng et al., 2009; Schumacher et al., 2017; MacCready et al., 2018).

To test if direct DNA binding stimulates the ATPase activity of McdA, we mutated a residue near its C-terminus identified by AF2 modeling and thermodynamic calculations (Pulianmackal et al., 2023). This His-SUMO-McdA[R158E] mutant forms an ATP-dependent dimer like WT McdA (Supplementary Fig. 6a), but with reduced DNA binding: it moderately shifted nsDNA and only at high protein concentrations (Supplementary Fig. 6b), highlighting the importance of this residue for DNA binding (Hester et al., 2007). His-SUMO-McdA[R158E] also has enhanced ATPase activity in the presence of DNA, although with specific activity of 21.8 hr^−1^, which is lower than that of WT McdA (Figure 3d), showing a direct correlation between DNA binding affinity and ATPase activity. Collectively, our data show that McdA non-specifically binds to DNA upon ATP binding and subsequent dimerization, consistent with putative models of the biochemical cycle of ParA ATPases (Leonard et al., 2005; Ptacin et al., 2010; Lim et al., 2014; Schumacher et al., 2017). Intriguingly, the ATPase activity of α-McdA is greatly stimulated by binding to DNA, suggesting that DNA binding may facilitate a conformational change in α-McdA that switches it to an ATP hydrolysis competent form.

### McdB stimulates the release of McdA from DNA without enhancing its ATPase activity in vitro

The activity of ParA ATPases is typically synergistically stimulated by their partner protein in the presence DNA (Davis et al., 1992; Ah-Seng et al., 2009; Schofield et al., 2010); therefore, we tested whether the ATPase activity of McdA was further enhanced by its partner protein, McdB. Surprisingly, McdB lowers His-SUMO-McdA ATPase activity (Figure 4a). Because McdB is concentrated at the carboxysomes (MacCready et al., 2018; Basalla et al., 2024), we tested whether high McdB concentrations (up to a 40:1 molar ratio) are needed for stimulation but found an McdB-dependent ATPase decrease (Supplementary Fig. 7).

**Figure 4:**
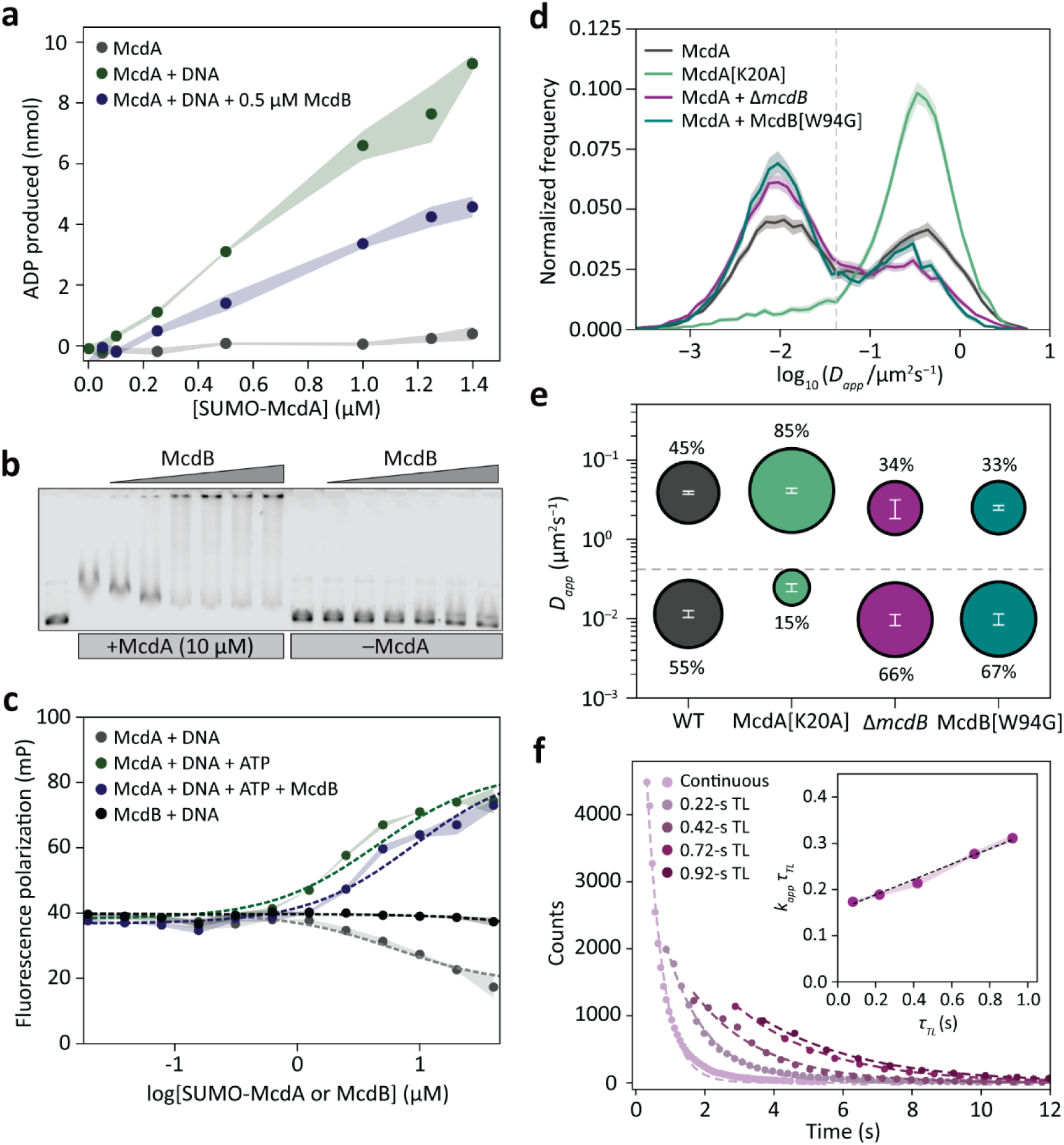
Carboxysome-localized McdB promotes release of McdA from DNA despite not stimulating its ATPase activity. **a.** ATPase activity of His-SUMO-McdA with DNA (0.1 mg/mL) and McdB (0.5 µM). His-SUMO-McdA with and without DNA is the same as Figure 3c. ATP concentration was kept fixed at 1 mM. Amount of ADP produced in reaction after 2 hr. Shading indicates the SEM of three independent experiments. **b.** EMSA with increasing concentrations of McdB (0.5, 1, 2.5, 5, 10, 12.5 µM) in the presence and absence of McdA (10 µM). First lane is DNA only. Second lane is DNA + McdA (10 µM). 1 mM ATP was present in all lanes. **c.** Binding curves of His-SUMO-McdA with DNA in the presence and absence of McdB (1 µM) probed by fluorescence polarization. Dashed lines are fits to a hyperbolic binding model. Shading indicates the SD of three independent experiments. **d.** Distributions of the apparent diffusion coefficient, *D_app_*, of McdA-PAmCherry^SW^ in WT cells (15,891 tracks from 335 cells over 4 independent experiments), McdA[K20A]-PAmCherrySW in WT cells (11,758 tracks from 149 cells over 3 independent experiments), McdA-PAmCherry^SW^ in Δ*mcdB* cells (20,097 tracks from 531 cells over 5 independent experiments), and McdA-PAmCherry^SW^ in McdB[W94G] cells (9,891 tracks from 480 cells over 2 independent experiments). Only tracks with ≥ 5 localizations are included. Lines and shaded regions indicate the mean ± 95% CI from 100 bootstraps. The dashed gray line is the theoretical lower bound for measurable *D_app_* based on the localization uncertainty from a fixed sample (Supplementary Fig. 9b). **e.** Bubble plot showing the apparent diffusion coefficients and associated population percentages for each of the two states in **d** from the fit to a Gaussian Mixture. Error bars are the standard deviations from independent experiments. **f.** Raw survival of fluorescence ‘on’ times for McdA-PAmCherry^SW^ in Δ*mcdB* cells across different time-lapse conditions, *τ_TL_*. Continuous imaging: *N* = 4487 tracks; *τ_TL_* = 0.22-s: *N* = 1990 tracks; *τ_TL_* = 0.42 s: *N* = 1227 tracks, *τ_TL_* = 0.72 s: *N* = 1139 tracks; *τ_TL_* = 0.92 s: *N* = 924 tracks). Dashed lines are fits by an exponential decay model with one apparent rate constant (see Methods). Inset: Product of the apparent rate constant, *k_app_*, and the time-lapse condition, *τ_TL_*, vs. *τ_TL_*. Shading is the error on the fit of the exponential decay model. Dashed line is the linear fit used to extract *k_diss_*.

Since DNA stimulates the ATPase activity of McdA, we tested whether McdB affects the binding of McdA to DNA. Using an EMSA with His-SUMO-McdA and McdB, we observed that, at low McdB concentrations (< 1:4 McdB:McdA molar ratios), McdB releases McdA from DNA (Figure 4b). At higher McdB concentrations (> 1:2 McdB:McdA molar ratios), we detected a higher-order nucleoprotein complex in the wells, which we interpret as a DNA-McdA-McdB complex (Figure 4b). McdB does not associate with the DNA substrate in the absence of McdA (Figure 4b). To measure the kinetics of McdA binding to DNA in the presence of McdB, we monitored the fluorescence polarization of a fluorescently labeled primer dimer (Figure 4c). In the presence of ATP, McdA readily binds to DNA with an equilibrium dissociation constant, *K*_d_, of 4.47 ± 0.01 µM. The addition of McdB increases *K*_d_ to 8.83 ± 0.87 µM, indicating that McdB reduces the binding affinity of McdA to DNA. This effect may be due to McdB promoting McdA release from DNA or inhibiting the binding of McdA to DNA. In either case, a decreased DNA-binding affinity may explain the reduced ATPase activity of McdA, as they are positively correlated (Figure 3).

We conclude that, unlike typical ParA/MinD systems (Davis et al., 1992; Ah-Seng et al., 2009; Schofield et al., 2010), the partner protein McdB does not enhance the ATPase activity of McdA but does interact with McdA to facilitate its release from DNA. This result contrasts with typical ParA systems, wherein partner proteins greatly stimulate ParA ATPase activity, which is thought, although not directly shown, to be directly coupled to release from the nucleoid (Davis et al., 1992; Ah-seng et al., 2009; Schofield et al., 2010; Vecchiarelli et al., 2013; Lim et al., 2014; Schumacher et al., 2017). Instead, McdA reaches similar ATPase activity upon DNA binding alone, and McdB facilitates McdA release without further stimulating its ATPase activity.

### Carboxysome-localized McdB facilitates McdA release from the nucleoid

Considering the effect of McdB on McdA complex assembly and release from DNA *in vitro* (Figure 4b), we explored whether these activities are preserved *in vivo*. Although bulk imaging data (Figure 2a-b) cannot detect McdA-McdB associations, we cannot exclude their contribution. To resolve the effects of this interaction with improved sensitivity, we measured the dynamics of McdA at a higher spatial-temporal resolution using live-cell PALM and single-molecule tracking (Manley et al., 2008) in *H. neapolitanus*. A two-state Gaussian Mixture Model fit well to the *D_app_* distribution of McdA-PAmC^SW^ trajectories from WT cells (Figure 4d-e, Supplementary Fig. 8, Table S1). The two components comprise 55 ± 7% and 45 ± 7% of the population, respectively: an immobile state (*D_app_* = 0.011 ± 0.001 µm^2^s^−1^) that is slower than our theoretical lower bound of detection (*D_app_* = 0.042 µm^2^s^−1^; Supplementary Fig. 9a), and a mobile state (*D_app_* = 0.387 ± 0.02 µm^2^s^−1^), which we assigned to free McdA diffusion.

To determine whether the slow state indicates α-McdA binding to DNA, we tracked α-McdA[K20A]-PAmC^SW^, the monomeric mutant with perturbed DNA binding, expecting a significant reduction in the slow state population. Fitting the *D_app_* distribution of α-McdA[K20A]-PAmC^SW^ results in two states comprising 85 ± 3% and 15 ± 3% of the population, respectively (Figure 4d-e, Supplementary Fig. 8, and Table S1): a dominant fast state with similar diffusion to McdA-PAmC^SW^, *D_app_* = 0.411 ± 0.03 µm^2^s^−1^, and a slow state (*D_app_* = 0.025 ± 0.003 µm^2^s^−1^). This shift in the *D_app_* distribution indicates an increase in the free population of McdA[K20A]-PAmC^SW^, consistent with its *in vitro* DNA-binding deficiency (Figure 2a-b and Figure 3b).

To measure the effect of McdB on McdA dynamics, we tracked McdA-PAmC^SW^ in Δ*mcdB* cells. Similar to WT cells, McdA-PAmC^SW^ in Δ*mcdB* cells shows two states with diffusion coefficients, 0.009 ± 0.002 µm^2^s^−1^ and 0.248 ± 0.07 µm^2^s^−1^ (Figure 4d-e, Table S1). However, the immobile population increases to 66 ± 6 %, indicating a ∼20% enrichment of DNA-bound McdA-PAmC^SW^ molecules in the absence of McdB, consistent with *in vitro* results showing that McdB promotes His-SUMO-McdA release from DNA (Figure 4b). Since most McdB molecules are carboxysome-associated (Basalla et al., 2024), we investigated whether the spatial distribution of McdB influences the dynamics of McdA. We tracked McdA-PAmC^SW^ in the presence of McdB[W94G], a mutant that does not associate with carboxysomes and is diffuse in the cytoplasm (Basalla et al., 2024). Since the N-terminus of McdB is predicted to associate with McdA (Pulianmackal et al., 2023) and the W94G mutation is at the C-terminus of McdB (Basalla et al., 2023), we hypothesized that McdB[W94G] would still interact with McdA. This mutation results in an McdA diffusion profile similar to that in Δ*mcdB* cells (Figure 4d-e, Supplementary Fig. 8, Table S1), showing that, under physiological conditions, McdB needs to be localized at carboxysomes to promote the release of McdA from DNA. It is also possible that, upon interacting with carboxysomes, McdB adopts a different conformation that enables its activity (Pulianmackal et al., 2023).

Given the high ATPase activity of McdA in the presence of DNA (Figure 3), we posited that McdA could also release from the nucleoid in an McdB-independent manner. To measure the *in vivo* basal dissociation rate, *k_diss_*, of McdA from DNA, we performed single-molecule time-lapse microscopy (see methods). We measured an α-McdA-PAmC^SW^ residence time of 6.02 ± 0.24 s and a dissociation rate of *k_diss_* = 0.166 ± 0.007 s^−1^ (Figure 4f). Strikingly, the rate of DNA release by α-McdA-PAmC^SW^ is 15-fold faster than the ATP turnover rate of His-SUMO-α-McdA (0.011 s^−1^) (Figure 3c-d). This result further supports that DNA release by α-McdA is not obligatorily coupled to ATP hydrolysis, consistent with increased His-SUMO-α-McdA release from DNA in the presence of α-McdB without enhanced ATP hydrolysis. Together, our data indicate McdA can release from DNA without McdB, but carboxysome-localized McdB facilitates this process.

### McdA dimers diffuse while bound to the nucleoid and turn over independently of McdB

In our Brownian ratchet model, ParA dimers can diffuse laterally along the nucleoid through transient unbinding and rebinding of DNA (Figure 5a) (Hu et al., 2017; Hu et al., 2021). In separate models for ParA-mediated cargo partitioning (Schumacher et al., 2017; Kohler et al. 2022), this property was proposed as a means to regulate the ParA dimer flux around the cargo, ultimately dictating the cargo motion. To determine if nucleoid-bound McdA dimers can diffuse beyond the elastic fluctuations of the chromosome (Figure 5a) (Lim et al., 2014; Surovtsev et al., 2016b), we extended our single-molecule imaging camera integration time to 80 ms, biasing our detection to only slower McdA-PAmC^SW^ molecules. Plotting the trajectory MSD versus time lag for nucleoid-bound McdA-PAmC^SW^ showed trajectories with heterogeneous behavior (Supplementary Fig. 10a). For this reason, we extracted diffusion coefficients from the cumulative distribution function (CDF) of McdA-PAmC^SW^ dimer squared displacements (*τ* = 80 ms; Supplementary Fig. 10b) (Schutz et al., 1997). The CDF is best fit by a two-state model, indicating two dynamic states for nucleoid-bound McdA-PAmC^SW^. Molecules in the slower state are immobile based on our measurable lower bound (*D_app_* < 0.0045 µm^2^s^−1^; Supplementary Fig. 9b), while the faster mobile state suggests lateral McdA diffusion on the nucleoid.

**Figure 5:**
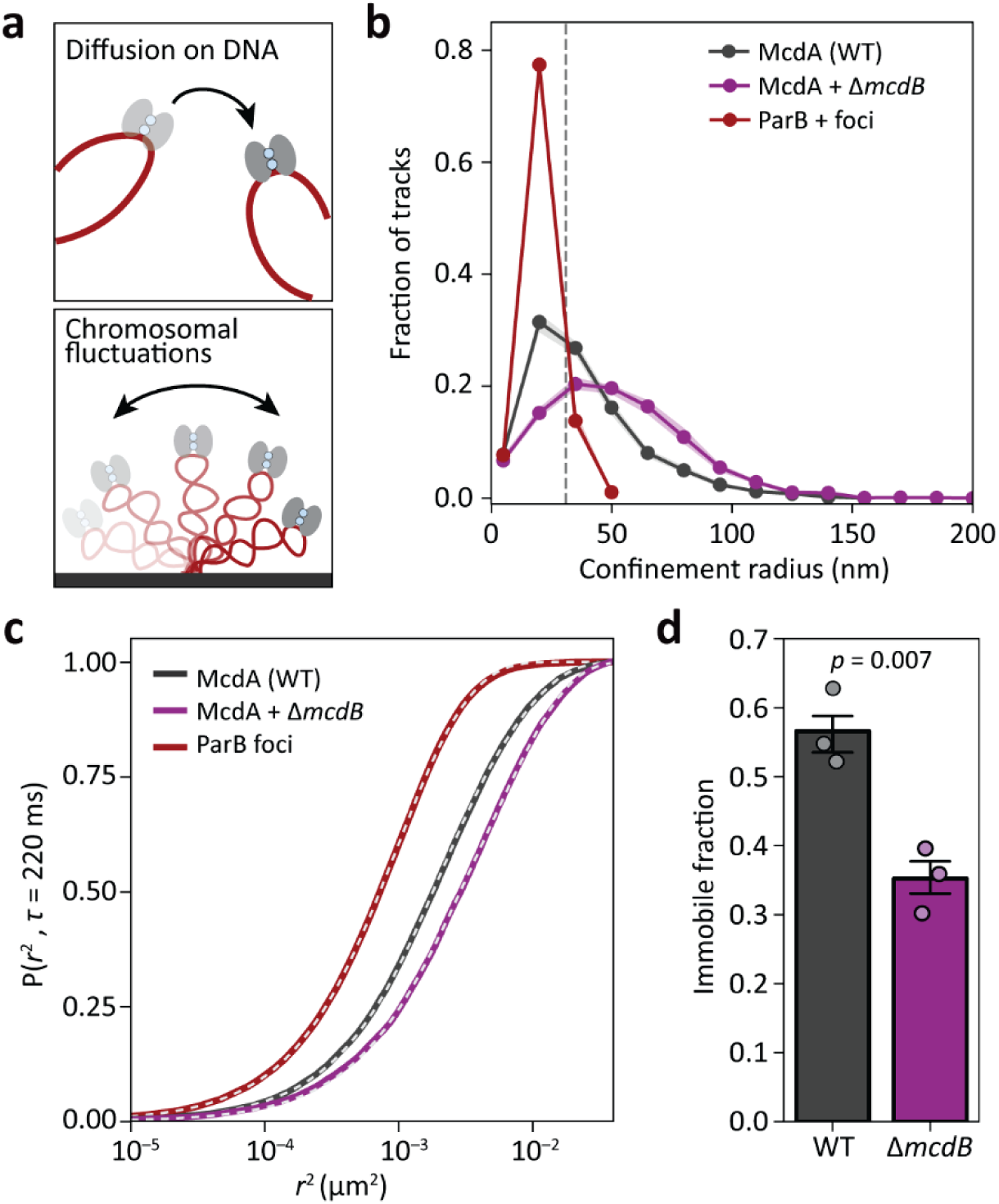
McdA dimers diffuse on the nucleoid. **a.** Possible mechanisms for the motion of nucleoid-bound McdA dimers (red): McdA dimers may hop from one DNA strand (gray) to another (top) or be limited to the elastic fluctuations of the chromosomal DNA (bottom). **b.** Confinement radius distributions for McdA-PAmCherry^SW^ in WT (*N* = 2366 tracks) and Δ*mcdB* (*N* = 2054 tracks) cells and ParB-mNG (*N* = 950 tracks). Solid lines and shaded regions indicate the mean ± 95% CI from 100 bootstrapped samples of the pooled datasets from 3 independent experiments. McdA-PAmCherry^SW^ tracks with confinement radii up to 0.031 µm (dashed gray line) are considered immobile. **c.** Cumulative distribution function, *P*(*r*^2^), of squared displacements from three pooled datasets of each strain. Dashed line is the two-state model fit (see Methods). **d.** Bar plot showing the immobile fraction (*D_app_* < 0.0016 µm^2^s^−1^) of McdA-PAmCherry^SW^ in WT and Δ*mcdB* cells. Data points are the fit results from independent experiments, and the error bars indicate the standard deviation across the experiments. *p*-value is from a Welch’s t-test.

To explain the diffusive states of nucleoid-associated McdA, we extended the observation period by adding a 140-ms dark time delay between imaging frames, and we tracked McdA-PAmC^SW^ as well as ParB-mNG foci as a chromosome marker. We found that the confinement radius of McdA-PAmC^SW^ in WT (47.0 ± 3.8 nm) and Δ*mcdB* (57.7 ± 4.7 nm) cells is greater than that of ParB-mNG foci (28.5 ± 0.9 nm), indicating that nucleoid-bound McdA is more mobile than the chromosome itself (Figure 5b, Table S2). Indeed, the mobile state of nucleoid-bound McdA-PAmC^SW^ (*D_app_*= 5.7 ± 0.6 × 10^−3^ µm^2^s^−1^) is more dynamic than that of ParB-mNG foci (*D_app_* = 1.5 ± 0.2 × 10^−3^ µm^2^s^−1^) (Figure 5c, Table S3), suggesting that McdA dimer diffusion results in nucleoid exploration length scales that are greater than the range of chromosomal fluctuations. Moreover, the *D_app_* of ParB-mNG foci is similar to that of the immobile state of McdA-PAmC^SW,^ implying that this population corresponds to McdA-PAmC^SW^ stably bound to a DNA strand. Notably, the immobile fraction of McdA-PAmC^SW^ is enriched in WT cells compared to Δ*mcdB* cells (Figure 5d, Table S3), suggesting that McdA diffusion on the nucleoid is inhibited by interactions with carboxysome-localized McdB.

### The mechanochemical Brownian ratchet model captures the distribution and dynamics of carboxysomes in H. neapolitanus

To gain insight into how the McdAB system mediates the partitioning of a large number of carboxysomes in *H. neapolitanus*, we built upon our established Brownian ratchet model in which the thermal motion of carboxysomes is rectified by a self-generated McdA gradient (MacCready et al., 2018; Byrne et al., 2025). This model is based on the established ParAB*S*-mediated low-copy plasmid partitioning model (Hu et al., 2015; Hu et al., 2017; Hu et al., 2021), with the parameters specified for and constrained by the carboxysome system in *H. neapolitanus*, as measured experimentally (Table S4).

Briefly, this computational model (Figure 6a) simulates the McdAB-mediated movement of carboxysomes (blue disks), on a flat 2D nucleoid surface (red). McdA-ATP dimers can ① bind to unoccupied spaces on this plane and ② diffuse laterally with a diffusion constant of 0.01 µm^2^s^−1^. McdB molecules are fixed on the carboxysome surface. Once McdA dimers bind to carboxysome-bound McdB, ③ they can no longer diffuse. The formed McdA-McdB bonds are modeled with chemical on and off rates and as elastic springs subject to thermal fluctuations, which stretch the bond and generate forces that drive carboxysome motion. The sum of the elastic forces from all of the McdA-McdB bonds on the surface of the carboxysomes yields a net force that drives the motion of the carboxysomes, which in turn breaks some of the old bonds and facilitates the formation of new ones. Importantly, ④ McdB accelerates the dissociation of McdA from the nucleoid. McdA can also ⑤ dissociate from the nucleoid independent of McdB. The dissociated McdA (McdA* in Figure 6a) must then ⑥ undergo nucleotide exchange and dimerization, resulting in a tens of seconds to minutes time delay before it can rebind to the nucleoid (Vecchiarelli et al., 2013). This delay creates an McdA-depletion zone behind the moving carboxysomes, which can be refilled either by cytosolic McdA dimers or through the lateral diffusion of nucleoid-associated McdA.

**Figure 6:**
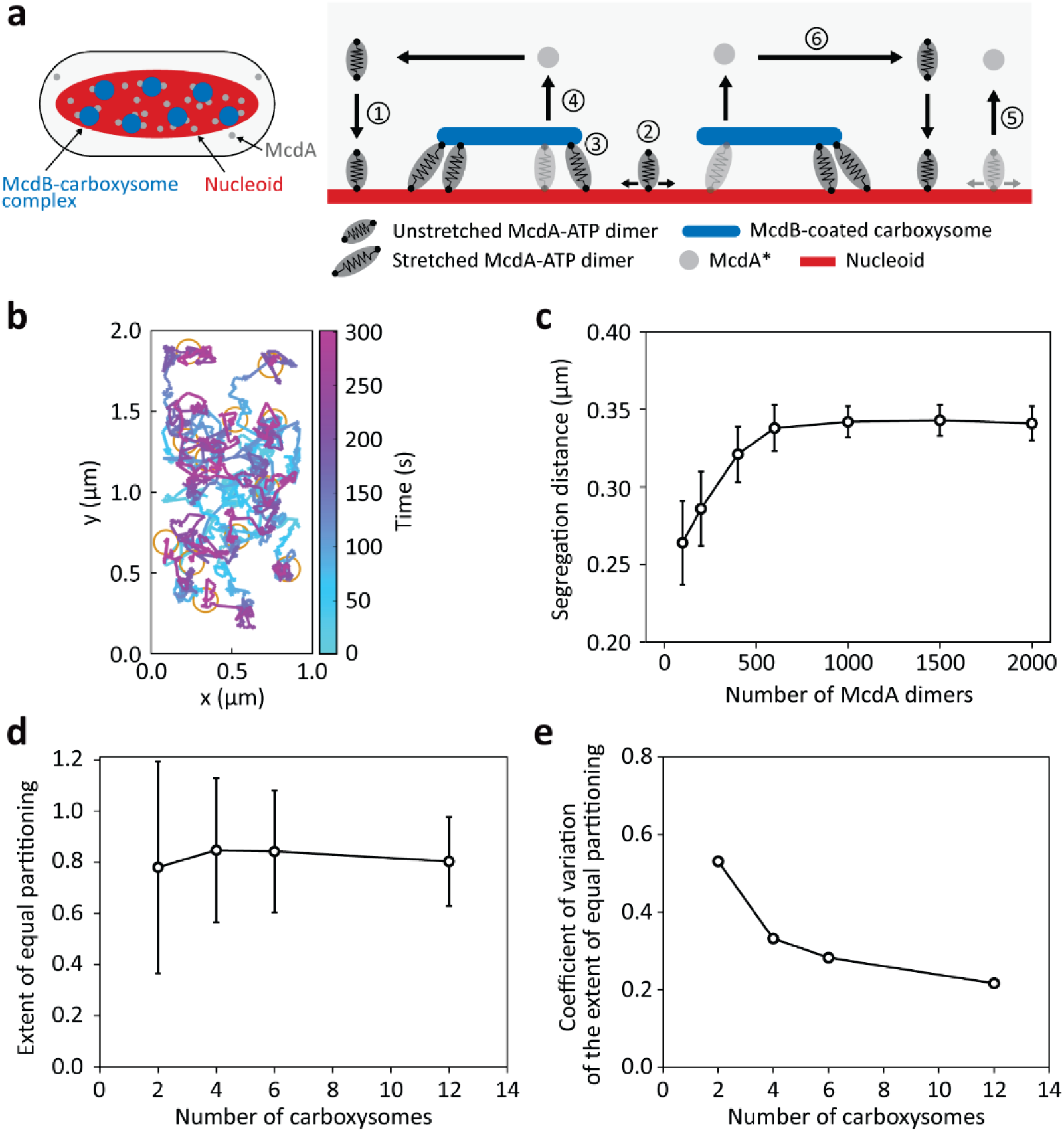
A Brownian ratchet model contextualizes carboxysome distribution by McdA. **a.** Schematic model of carboxysome distribution. Left: Key components in the carboxysome positioning system. Right: Brownian ratchet model for carboxysome distribution. See main text for details. **b.** Example simulated 12-carboxysome system showing the trajectories of the carboxysomes. The carboxysomes are initially clustered at the middle of the simulation region, and their final positions at the end of the simulation are denoted by the yellow circles. **c.** Average carboxysome segregation distance as a function of McdA amount in the simulation. Error bars denote the standard deviation from 100 simulations. **d.** Extent of equal partitioning as a function of carboxysome number in the simulation. Error bars denote the standard deviation from 100 simulations. **e.** Variation associated with the extent of equal partitioning as a function of carboxysome number.

We first simulated the partitioning of twelve initially clustered carboxysomes by 200 McdA dimers within a 1 µm × 2 µm simulation box (Hu et al., 2017). In the first half of the 300-s simulation, carboxysomes exhibit biased random motion away from one another (Figure 6b), resulting in an average segregation distance of approximately 0.28 µm (Figure 6c). This result is consistent with our experimentally measured nearest-neighbor carboxysome spacing of approximately 0.3 µm. In the second half of the 300-s simulation, carboxysomes exhibited constrained motion, resembling our experimental results (Figure 1). Given that previous reports indicate ParA ATPase abundances ranging from 80 to 2300 dimers (Bouet et al., 2005; Adachi et al., 2006; Lim et al., 2014; Lioy et al., 2015; Schumacher et al., 2017), we examined how McdA copy number influences carboxysome dynamics. We found a positive correlation between the number of McdA dimers and the carboxysome segregation distance, which plateaus at approximately 0.33 µm beyond 500 dimers (Figure 6c). This result indicates that a few hundred dimers of McdA are sufficient to efficiently distribute a large number of carboxysomes.

Considering that carboxysomes are present in much higher copy numbers than most cargos positioned by ParA ATPases, we evaluated how the McdA-specific parameters perform when partitioning a smaller number of carboxysomes. We calculated the extent of equal partitioning, defined as the weighted probability of achieving an equal partition: for a system with 12 carboxysomes, a 6/6 partitioning contributes 1 to this quantity, a 4/8 partitioning contributes 0.5, and so on. Our findings indicate that high partitioning fidelity is maintained in simulations involving 2 – 12 carboxysomes (Figure 6d). However, the variability in fidelity decreases as carboxysome number increases (Figure 6e), suggesting that a greater number of cargos in the system averages out stochastic fluctuations and enhances the robustness of partitioning. Importantly, these results highlight that the McdAB system does not necessarily ensure equal partitioning of carboxysomes between the cell halves; instead, it prevents carboxysomes from aggregating.

## DISCUSSION

In this study, we characterized the dynamics of α-carboxysomes and the properties of their positioning system, α-McdAB, using complementary *in vitro*, *in vivo*, and *in silico* approaches. Our results indicate that several aspects of this system are consistent with a Brownian ratchet mechanism (Hu et al., 2015; Hu et al., 2017; Hu et al., 2021). First, the distribution of McdA on the nucleoid is asymmetric in nature (Figure 2), satisfying a requirement for a ‘ratchet’-based mechanism. Second, McdA forms ATP-dependent dimers that enable its association with DNA (Figure 3), consistent with the initial biochemical model steps. Third, carboxysome-localized McdB facilitates the release of McdA from the nucleoid (Figure 4), creating an McdA depletion zone around carboxysomes. Fourth, McdA dimers can diffuse while bound to the nucleoid (Figure 5), providing a mechanism to replenish the McdA depletion zones. Finally, the increased immobility of nucleoid-bound McdA dimers in the presence of McdB suggests that the carboxysome-McdB-McdA-DNA complex acts as a diffusion-restricting tether. Based on these observations, we conclude that α-McdAB employs a Brownian ratchet mechanism to distribute carboxysomes in *H. neapolitanus*.

### Diffusion of McdA/ParA dimers on the nucleoid regulates cargo dynamics

Our finding that α-McdA dimers can diffuse on the nucleoid indicates a characteristic shared by other ParA ATPases *in vitro* and *in vivo* (Kiekebusch et al., 2012; Hwang et al., 2013; Vecchiarelli et al., 2013; Surovtsev et al., 2016b; Schumacher et al., 2017). In *Caulobacter crescentus*, intermittent DNA binding by chromosome-segregating ParA establishes the gradient necessary for directional transport (Surovtsev et al., 2016b). Similarly, ATPases like PomZ and MipZ, which position division machinery in *Myxococcus xanthus* and *C. crescentus*, respectively, are mobile even in their ATP hydrolysis-incompetent forms (Kiekebusch et al., 2012; Schumacher et al., 2017), supporting ATP-hydrolysis-independent diffusion on the nucleoid. Furthermore, single SopA-GFP molecules show intersegmental hopping on DNA carpets in flow cells, though at a faster rate (∼0.9 µm^2^s^−1^) than in living cells (Vecchiarelli et al., 2013). This discrepancy is likely due to the lower DNA concentration in flow cells, which allows SopA-GFP dimers to diffuse over longer distances before rebinding DNA, an effect that would be limited in the crowded cellular environment.

The prevalence of nucleoid-bound diffusion by ParA ATPases highlights its role in regulating cargo dynamics. This property is essential for a ‘flux balance’ mechanism, which has been proposed as a mechanism for ‘true’ or regular cargo positioning (Ietswaart et al., 2014; Schumacher et al., 2017; Murray and Howard, 2019; Kohler et al., 2022). In regular positioning, cargo motion is biased towards specific or ‘true’ locations on the nucleoid in a cargo copy-dependent manner (mid or quarter nucleoid positions for one or two cargos, respectively). Cargos move in the direction of the greatest incoming ATPase flux until they reach their ‘true’ position, where the fluxes around the cargo are balanced. For a cargo to receive positional information through these ATPase fluxes, the nucleoid-bound ATPase must diffuse sufficiently to interact with the cargo. In a single cargo system, this means that the ATPase must be able to diffuse at least half the length of the nucleoid. In the case of the ParA homolog PomXYZ, which positions the *M. xanthus* divisome, the imbalance in the flux of dimers of PomZ (the ParA ATPase) on either side of the PomXY cluster (the cargo) drives the PomXY cluster towards mid-cell, where PomZ fluxes are balanced (Schumacher et al., 2017).

When the cargo copy number is drastically increased—like in the case of carboxysomes, which can be present in an excess of a dozen copies—the necessary diffusive length scale for McdA dimers decreases because each molecule is more likely to encounter a carboxysome (Azaldegui et al., 2025). Supporting this notion, our measurements indicate that the average diffusive length scale of α-McdA dimers is 0.26 µm, while the average distance between carboxysomes is approximately 0.30 µm (Supplementary Fig. 1). Consequently, α-McdA dimers are always able to interact with a carboxysome, resulting in our experimental and computational observations of tight local excursions.

### McdA release from DNA is not obligatorily coupled to ATP hydrolysis

In classical ParA ATPase models, the release of ParA dimers from DNA is proposed to be tightly coupled to ParB-stimulated ATP hydrolysis (Leonard et al., 2005; Ptacin et al., 2010). ATP hydrolysis destabilizes the dimers, causing their release as monomers. Stochastic models use these hydrolysis rates to model ParA dissociation kinetics from DNA (Lim et al., 2014; Hu et al., 2015; Köhler et al., 2022). However, our data contradict this model: the α-McdA dissociation rate (0.166 ± 0.007 s^−1^) is ∼15-fold faster than its ATP hydrolysis rate (0.01 s^−1^), and α-McdB stimulates α-McdA dissociation without enhancing ATP hydrolysis. This accelerated DNA dissociation relative to ATP hydrolysis is also seen in the partitioning ATPases SopA of the F plasmid and ParA of the P1 plasmid (Hwang et al., 2013; Vecchiarelli et al., 2013). Importantly, while ATP hydrolysis does result in release from DNA, we propose these activities are not obligatorily coupled—release from DNA can occur independently of ATP hydrolysis.

### Comparing the α- and β-McdAB systems

Our mechanochemical Brownian ratchet model effectively captures the observed cargo dynamics in both the α- and β-McdAB, despite several key differences between the two systems. For instance, β-McdA forms oscillating gradients that traverse the entire *S. elongatus* nucleoid every 15 minutes (MacCready et al., 2018; Hakim et al., 2021; Dudley et al., 2025). This behavior contrasts sharply with the short-scale, rapidly rearranging gradients formed by α-McdA. This difference may stem from the distinct putative ATP-binding pockets and associated ATPase activities between the two McdA proteins. Notably, β-McdA lacks the conserved ‘signature’ lysine in the Walker A motif and exhibits high ATPase activity even in the absence of DNA and McdB (MacCready et al., 2018; Hakim et al., 2021). The presence of McdB further stimulates this activity approximately 2-fold (MacCready et al., 2018). In contrast, α-McdA alone shows negligible ATPase activity, which is drastically enhanced only in the presence of DNA and is not further stimulated by McdB. Despite these differences in ATPase activity, the McdB protein of both systems can displace McdA from DNA, fulfilling the requirements of our Brownian ratchet model.

Another distinction between the two systems is that β-carboxysomes in *S. elongatus* are generally less abundant than α-carboxysomes in *H. neapolitanus*; specifically, there are fewer carboxysomes per micron of cell length (∼1.4 vs 2.7 foci per micron) (Hakim et al., 2021; Dudley et al., 2025). This reduced carboxysome density leads to larger spacing between carboxysomes: 0.6 µm for β-carboxysomes compared to 0.3 µm for α-carboxysomes. Despite these differences, the product of focus density and spacing is consistent across both systems, maintaining an approximate value of 0.8. Additionally, both α- and β-carboxysomes exhibit similar diffusive dynamics; both types have a radius of confinement of approximately 40 nm (Dudley et al., 2025). These tight local excursions are also indicative of ‘true’ positioning, suggesting a conserved mechanism across the two systems that is regulated by the interaction kinetics and subsequent ATPase diffusion.

In conclusion, this study has uncovered key properties of the McdAB system that is responsible for positioning α-carboxysomes. Our data demonstrate that the dynamics of α-carboxysomes and α-McdA align with a Brownian ratchet mechanism. Crucial to this finding was the integration of *in vitro* biochemical assays and computational modeling with live-cell single-molecule imaging, showcasing the effectiveness of these complementary approaches in elucidating molecular mechanisms in bacteria. We anticipate that our results for α-carboxysomes will be relevant to understanding the spatial organization of other bacterial microcompartments.

## Materials and Methods

### Construct designs and cloning

Strains, plasmids, and oligos used in this study are listed in Tables S5, S6, and S7, respectively. All constructs were generated using Gibson Assembly or site-directed mutagenesis (SDM) (NewEngland Biolabs Q5 Hot Start High-Fidelity; Cat. #M0494S) and verified by sequencing. Fragments for assembly were synthesized by polymerase chain reaction (PCR) or ordered as gBlocks (IDT). Constructs contained flanking DNA that ranged from 750 to 1100 bp in length upstream and downstream of the targeted insertion site to promote homologous recombination into the target genomic locus. Cloning of plasmids was performed in chemically competent *E. coli* Top 10 or Stellar cells (Takara Bio).

To prevent phenotype recovery by the presence of the native unlabeled McdA, all *mcdA* fusion plasmids were transformed into an *H. neapolitanus* Δ*mcdA* strain (AGV_Hn100) (MacCready and Tran et al., 2021) or a Δ*mcdAB* strain (AGV_Hn432), unless otherwise noted. The mutants were selected by plating on S-6 agar plates supplemented with the appropriate antibiotic. All fusions were verified by PCR and by visualization of fluorescence signal by microscopy.

The Δ*mcdAB* background strain (AGV_Hn432) was used to transform fluorescent fusions of McdA for complementation. Strain AGV_Hn432 was constructed as follows. The sequence encoding plasmid pLT90 was amplified by PCR from pLT104 (MacCready and Tran et al., 2021) using the primer pair Pterin_rev/Hn0910_fwd. This step removed the *mcdB* gene (*Hn0911*) from pLT104. pLT90 was then transformed into AGV_Hn2 and selected by spectinomycin resistance to generate AGV_Hn432.

The strain AGV_Hn321 was used to induce the expression of McdB-PAmCherry and determine the localization uncertainty of PAmCherry molecules in fixed *H. neapolitanus*. This strain was constructed by amplifying a fragment from pLT17 (MacCready and Tran et al., 2021) using primer pair ptrc_backbone_rev/mcdB_CO_fwd and the *PAmCherry* fragment with primer pair PAmCherry_fwd/rev. The plasmid pLT67 was produced by Gibson assembly of the two fragments. pLT67 was then transformed into AGV_Hn2 and selected by kanamycin resistance to generate AGV_Hn321.

The strains AGV_Hn404 and AGV_Hn518 are native fluorescent fusions of McdA at its N- and C-terminus, respectively. The plasmid encoding *PAmCherry-mcdA* (pLT79) was produced by Gibson assembly of the fragment *PAmCherry-mcdA* amplified by primer pair PAmCherry_fwd/mcdA_CO_rev with the backbone amplified by primer pair Hn_0913_rev/mcdA-mcdB-CO_fwd. The plasmid encoding *mcdA-PAmCherry* (pLT87) was produced by Gibson assembly of the *PAmCherry* fragment amplified by primer pair GSGx9-PAmCherry_fwd/pamcherry_rev_2 with the backbone amplified by primer pair PAmCherry-mcdB/GSGx9_mcdA_rev from pLT79. Both plasmids were transformed into AGV_Hn2. The mutants were selected by plating on S-6 agar plates supplemented with 25 µg/mL of chloramphenicol.

The native sandwich fusions of McdA in strains AGV_Hn532 (mNeonGreen) and AGV_Hn431 (PAmCherry) were used for bulk imaging and single-molecule tracking of McdA, respectively. First, the sequence encoding a *PAmCherry* fragment with flanking GSGSGS linker sequences was amplified with primer pair GSGSGS_PAmCherry_fwd/rev. The *PAmCherry* fragment was then inserted in between the codon sequences encoding *mcdA* residues 59 and 60, amplified by primer pair GS_mcdA_fwd/Hn0912_GS_rev, by Gibson assembly to produce pLT94. The gene sequence for *mcdB* was included downstream of *mcdA*, followed by a chloramphenicol resistance cassette. The *cso* promoter was duplicated downstream of the resistance cassette to maintain faithful expression of the rest of the operon. To produce pCA2 (*mcdA-mNeonGreen^SW^*), the backbone from pLT94 was amplified using primer pair GS_mcdA_fwd_2/Hn0912_GS_rev and Gibson assembled with an *mNeonGreen* fragment amplified by primer pair GS-mNG_fwd/mNG-GS_rev. To prevent phenotype recovery by the presence of the native unlabeled McdA, *mcdA* fusion plasmids were transformed into a Δ*mcdA* strain (AGV_Hn100) (MacCready and Tran et al., 2021). The mutants were selected by chloramphenicol resistance.

To generate the Δ*mcdB* constructs, first, the plasmid LT103 was generated by amplifying pLT94 using the primer pair delmcdB_fwd/delmcdB_rev to remove the *mcdB* sequence. The amplified sequence was then ligated using the KLD Enzyme Mix (New England Biolabs). Plasmid pLT103 was then transformed into AGV_Hn432 to generate AGV_Hn441. To generate the strain AGV_Hn498, the backbone sequence of pLT103 was amplified using GS_mcdA_fwd_2/Hn0912_GS_rev and Gibson assembled with a *mNeonGreen* fragment amplified by primer pair GS-mNG_fwd/mNG-GS_rev. The resulting plasmid pCA3 was then transformed into AGV_432. The mutants were selected by chloramphenicol resistance.

Constructs containing point mutations were generated by Site-Directed Mutagenesis (SDM). To generate the McdA[K20A] constructs, the plasmids pCA2 and pLT94 containing the sandwich fusions for wildtype McdA with mNeonGreen and PAmCherry, respectively, were amplified using the primer pair SDM_McdA_K20A_fwd_long/mcdA_K20R_rev_long. The amplified sequences were then ligated using the KLD Enzyme Mix (New England Biolabs). The resulting plasmids pCA4 and pLT104 were then transformed into a Δ*mcdAB* strain (AGV_Hn432) to generate strains AGV_Hn481 (McdA[K20A]-mNG^SW^) and AGV_537 (McdA[K20A]-PAmC^SW^). The mutants were selected by chloramphenicol resistance.

To generate the McdB[W94G] construct, the plasmid pLT94 was amplified using the primer pair SDM_McdB_W94G_fwd/ SDM_McdB_W94G_rev. The amplified sequence was then ligated using the KLD Enzyme Mix (New England Biolabs). The resulting plasmid pCA31 was then transformed into AGV_432 to generate strain AGV_554.

### Media and growth conditions

All mutants described in this study were constructed using WT *Halothiobacillus neapolitanus* c2 (*H. neapolitanus*) (Parker) Kelly and Wood (ATCC 23641) purchased from ATCC. Cultures were grown in ATCC Medium 290: S-6 medium for *Thiobacilli* (Hutchinson et al., 1965) and incubated at 30 °C while shaken at 130 RPM in air supplemented with 5% CO_2_, unless otherwise stated. Strains were preserved at −80 °C in 10% DMSO.

### Making competent cells of *H. neapolitanus*

*H. neapolitanus* competent cells were produced as previously reported (Pulianmackal et al., 2023). Briefly, 0.5 L of culture was grown to an OD of 0.03 – 0.07. Cultures were harvested by centrifugation at 5000 ×g for 20 min at 4 °C. Pellets were resuspended and washed twice with 250 mL of ice-cold nanopore water. All wash centrifugation steps were performed at 3000 ×g for 30 min at 4 °C. The final pellet after washing was resuspended in 2 mL of ice-cold nanopore water. Competent cells were used immediately or frozen at −80 °C for future use. Frozen competent cells were thawed at 4 °C before use.

### Transformation in *H. neapolitanus*

50 – 100 µL of competent cells were mixed with 5 – 7 µL of plasmid DNA (0.5 – 3 µg) and incubated on ice for 5 min. The mixture of competent cells and DNA was then transferred to a tube containing 5 mL of ice-cold S-6 medium without antibiotics and incubated on ice for 5 min. Transformations were recovered for 18 – 22 h while shaken at 130 rpm at 30 °C in air supplemented with 5% CO_2_. Clones were selected by plating on S-6 plates with appropriate antibiotics. Colonies were restreaked once or twice. Restreaked colonies were verified for mutation by PCR and fluorescence microscopy.

### Protein structure prediction and visualization

Protein structures were predicted using the ColabFold (Mirdita et al., 2022) implementation of AlphaFold2 (Jumper et al., 2021). Molecular graphics and protein structure analyses were performed in PyMOL (Delano 2008).

### Expression and purification of WT and mutants of H. neapolitanus McdA proteins

Wildtype and mutant *H. neapolitanus* McdA proteins were expressed with an N-terminal His-SUMO tag in *E. coli* BL21(DE3) using a pET15b vector. The overnight culture was grown in LB media supplemented with 100 μg/mL carbenicillin. 1% of the overnight culture was transferred into 1 L of fresh LB media. Protein expression was induced with 100 μM IPTG when the OD600 of the culture reached 0.5-0.7. Cultures were incubated at 16 °C for 16 hours following induction. Cells were harvested by centrifugation, and pellets were stored at −80 °C. The cell pellet was resuspended in 30 mL of lysis buffer: 50 mM Tris, 1 M KCl pH 8, 10% glycerol, 5 mM β-mercaptoethanol (BME), 1 protease inhibitor tablet (Thermo-Fisher). 30 mg of lysozyme was added to the resuspended pellet, and the suspension was incubated on ice for 10 minutes. Cells were disrupted by sonication using 10-second pulses (on) followed by 20-second intervals (off) at 50% power for 10 minutes. The lysate was centrifuged at 15,000g for 30 minutes at 4 °C, and the supernatant was filtered through a 0.45 μm filter. The filtered lysate was loaded onto a 5 mL HisTrap HP column (Cytiva) pre-equilibrated with Buffer A (50 mM Tris, 1 M KCl pH 8, 10% glycerol, 5 mM BME). The column was washed with 5-column volumes of 5% Buffer B (50 mM Tris, 1 M KCl, 500 mM imidazole pH 8, 10% glycerol, 5 mM BME). The protein was eluted using a 5-100% gradient of Buffer B on an AKTA Pure system. Pooled fractions were injected onto a HiLoad 16/600 Superdex 200 pg column (GE Healthcare Life Sciences) equilibrated with Buffer A. Peak fractions were concentrated and filtered through a 0.45 μm filter and stored at −80 °C.

### Gel filtration assay

50-µL samples of SUMO-Hn McdA or its mutants (at approximately 50 µM) with and without 1 mM ATP were incubated at room temperature before injection into a Superdex 200 5/150 GL column (GE Healthcare Life Sciences). For reactions without ATP, the column was equilibrated with 50 mM Tris-HCl (pH 8), 1 M KCl, 10% glycerol, 5 mM β-mercaptoethanol (BME), 5 mM MgCl₂ and for reactions with ATP the column was equilibrated with 50 mM Tris-HCl (pH 8), 1 M KCl, 10% glycerol, 5 mM BME, 5 mM MgCl₂, 1 mM ATP. The flow rate was set to 0.17 mL/min, and elution profiles were monitored by measuring absorbance at 280 nm.

### DNA-binding assay

Electrophoretic mobility shift assays (EMSAs) were performed in a final reaction volume of 10 μL using 50 mM Tris, 1 M KCl pH 8, 10% glycerol, 5 mM BME, 5 mM MgCl_2_ buffer. The DNA substrate was a pUC19 plasmid (10 nM). SUMO-Hn McdA or its mutants were incubated with the DNA for 30 minutes at 23 °C in the presence of either 1 mM ATP or 1 mM ADP, at the indicated protein concentrations. Where applicable, McdB was added at the specified concentrations. Following incubation, 1 μL of DNA loading dye was added to each reaction. Samples were then loaded onto a 1% agarose gel prepared in TAE buffer with ethidium bromide and run at 100 V for 60 minutes. After electrophoresis, the gel was visualized using a Li-Cor Odyssey imager.

### ATPase assay

ATPase assays were performed in a buffer containing 50 mM Tris, 1 M KCl pH 8, 10% glycerol, 5 mM BME, 5 mM MgCl_2_, and 0.1 mg/mL sonicated salmon sperm DNA (unless otherwise specified). The reaction mixture, at the indicated protein concentration, was incubated at room temperature for 10 minutes. Following incubation, 10 μL of the reaction mixture was combined with 80 μL of EnzChek phosphate release assay (ThermoFisher Scientific) solution and further incubated for 5 minutes at room temperature. To initiate the ATPase reaction, 10 μL of 10 mM ATP was added to the mixture, which was then transferred to a 96-well plate. The plate was continuously shaken, and absorbance at 360 nm was measured every 2.5 minutes for 160 minutes using a TECAN Infinite M200 Pro plate reader. ATP hydrolysis was quantified by comparing the absorbance data to a phosphate standard curve. ATP turnover was determined by the linear regression of free phosphate produced vs. time up to 2 hours. Specific activities were determined by linear regression of ATP turnover (nmol h^−1^) vs. [McdA] nmol. ATP turnover linear regressions were weighted by the error (SD) of three replicates, and specific activity linear regressions were weighted by the error on fit from the ATP turnover linear regression.

### Fluorescence polarization assay

Two single-stranded oligonucleotides were purchased from IDT: one with a FAM fluorophore at the 5’ end (FAM-5’-GTGTGTGTGTGTGTGTGTGTGTGT-3’) and a complementary strand without a fluorophore. The oligonucleotides were dissolved in water to a concentration of 100 µM. Equal volumes of each primer were mixed, heated to 95 °C for 10 minutes, and gradually cooled to 72 °C. The reaction was then brought to room temperature by lowering the temperature by 1 °C per cycle. The resulting primer dimers were stored at −20 °C for further use. SUMO-McdA was exchanged into a buffer containing 50 mM Tris (pH 8), 200 mM KCl, 10% glycerol, and 5 mM BME. Fluorescence polarization experiments were performed using the same buffer, supplemented with 5 mM MgCl_2_, 1 mM ATP, and 100 nM DNA (primer dimers). Fluorescence polarization measurements were conducted in triplicate at the specified concentrations of SUMO-McdA using a TECAN Infinite M1000PRO plate reader. Excitation and emission wavelengths were set to 470 nm and 517 nm, respectively, with a bandwidth of 10 nm. polarization traces for each sample were fit to a hyperbolic binding model using a weighted least-squares regression. Errors equal to 0 were set to 0.01 to prevent division by zero.

### Cleaning coverslips for fluorescence microscopy

For widefield epifluorescence microscopy, 35-mm glass-bottomed dishes (MatTek, catalog number P35G-1.5-14-C) were rinsed with 70% ethanol and dried in an oven at 30 °C. For single-molecule imaging, large (35 × 50mm) and small (25 × 25 mm) coverslips (Fisherbrand) were plasma-etched in argon for 15 min.

### Widefield epifluorescence microscopy

All live-cell microscopy was performed using exponentially growing cells. 2 – 4 µL of cells were spotted onto a pad of 2% UltraPure agarose (Invitrogen, catalog number 16500) + S-6 and imaged on a 35-mm glass-bottomed dish (MatTek, catalog number P35G-1.5-14-C). All fluorescence and phase contrast imaging were performed using a Nikon Ti2-E motorized inverted microscope controlled by NIS Elements software. The microscope was equipped with a SOLA 365 LED light source, a 100× oil immersion objective lens (Oil CFI Plan Apochromat DM Lambda Series for Phase Contrast), and a Hamamatsu Orca Flash 4.0 LT + sCMOS camera. CbbS-mTurquoise2 (CbbS-mTQ) labeled carboxysomes were imaged using a “CFP” filter set (C-FL CFP, Hard Coat, High Signal-to-Noise, Zero Shift, Excitation: 436/20 nm [426-446 nm], Emission: 480/40 nm [460-500 nm], Dichroic Mirror: 455 nm). ParB-mNG, mNG-FliN, and McdA-mNG^SW^ were imaged using a ‘YFP’ filter set (C-FL YFP, Hard Coat, High Signal-to-Noise, Zero Shift, Excitation: 500/20 nm [490-510 nm], Emission: 535/30 nm [520-545 nm], Dichroic Mirror: 515 nm). DAPI-stained nucleoids were imaged using a ‘DAPI’ filter set (C-FL DAPI, Hard Coat, High Signal-to-Noise, Zero Shift, Excitation: 350/50 nm [325-375 nm], Emission: 460/50 nm [435-485 nm], Dichroic Mirror: 400 nm). Samples were imaged for 1-1.5 hr before loading a new sample.

### Image processing and analysis

Software packages used in this study are listed in Table S9 and available in the Biteen lab Github repository: https://github.com/BiteenMatlab/McdA_carboxysomes.

#### Cell registration

Image stacks were drift-corrected using a custom Python script: *cell_registration.py*. First, the phase contrast images were registered to the first frame in the stack using cross-correlation (*skimage.feature.register_translation* and *skimage.transform.warp*). If the stack contained multi-channel data, the shifts used for the phase contrast images were applied to all other time-corresponding frames in other channels. The registered stacks were then saved as separate TIFF stacks.

#### Cell segmentation

Drift-corrected phase contrast images were used for cell segmentation via the Omnipose (Cutler et al., 2022) package in Python. A custom Python script was implemented for batch segmentation: *PhaseMasks_omni.py*. Prior to segmentation, a Gaussian blur (SD of Gaussian = 1 pixel; *skimage.filters.gaussian*) was applied to the images. The *bact_phase* pre-trained model was used for segmentation. Cells touching the borders of the image were ignored, segmented objects smaller than 50 pixels were removed, and erroneous segmentations were manually corrected using the Omnipose GUI or excluded from further analysis if cell boundaries were not clear from the phase image. Resulting phase masks were used for subsequent single-cell analyses.

#### Carboxysome focus detection

To measure the number of foci per cell length, the cell length was first measured from the phase masks. Carboxysome foci were detected using a custom Python script: *detect_carboxysomes.py*. First, the phase masks were used to individually analyze every cell in the field of view. The fluorescence signal in each cell was then sharpened using the scikit-image (van der Walt et al., 2014) function *skimage.filters.unsharp_mask*; this step was followed by a Gaussian blur (*skimage.filters.gaussian*). Using this preprocessed image, carboxysome foci were detected with the Laplacian of Gaussian (LoG) algorithm (*skimage.feature.blob_log*). To reduce the effects of overlapping carboxysome PSFs and improve detection, analysis was performed on image stacks that captured carboxysome separation due to their motion. All preprocessing and detection were performed on all frames within a time-lapse image stack. The frame with the highest number of detected foci was used for further analysis. In this chosen frame, putative foci signals were fit to a 2D Gaussian for subpixel localization. Carboxysome foci with a standard deviation on the 2D Gaussian fit less than 1 pixel (66 nm) were excluded. Nearest neighbor distances between adjacent carboxysome foci were determined using the custom function *nearest_neighbor*. The number of carboxysome foci localized in a cell was normalized to the length of that cell.

#### Carboxysome and chromosome tracking

Time-lapse fluorescence microscopy was used to measure the motion of carboxysome foci and the chromosome (ParB-mNG foci). The signal was collected with 200-ms exposures and a dark-time delay of 800 ms. The image stack length was optimized for each sample: 10 frames for Δ*mcdA*, PAmCherry-McdA, and McdA-PAmCherry, and 25 – 31 frames for all other strains. Tracking of drift-corrected image stacks was done with the TrackMate (Tinevez et al., 2017) plugin in Fiji. Fluorescent spots were detected with LoG (radius = 3 pixels or 198 nm) using subpixel localization. Localizations were then linked with the LAP tracker method using a maximum linking length of 2 pixels (132 nm) and no gaps allowed. To remove erroneous trajectories due to mis-linking of proximal or overlapping foci, only trajectories with lengths equal to the full imaging collection period were used for MSD analysis. Each trajectory was analyzed independently to extract a trajectory-average apparent diffusion coefficient, *D_app_*, and the anomalous diffusion exponent *α*. The mean-squared displacement, 〈*r^2^*〉, as a function of time lag (*nτ*), where *n* is an integer, and *τ* is the 1-s time between frames, was calculated for up to the first 20 *τ*. The first 8 *τ* of the resulting curve were fit to Equation 1 using weighted non-linear least squares (*scipy.optimize.curve_fit*). The following bounds were placed on the fit parameters: *D_app_* [0, infinity]; *α* [0, 2]. The distribution *D_app_* was then fit to a Gaussian Mixture Model (GMM) (*sklearn.mixture.GaussianMixture*) with one or two components, which provides the average *D_app_*, the standard deviation, and the population weight associated with each component.

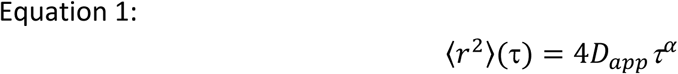

For velocity autocorrelation function (VACF) analysis, the same trajectory length thresholds described above were used. To calculate the VACF, the displacement, *r*, of a particle was used to calculate the velocity, *v*, after the time step, δ = 1s. We then calculated the dot product of the particle velocity and its velocity after some time lag, *τ*, as shown in Equation 2. The VACF was then normalized to *τ* (0 s).

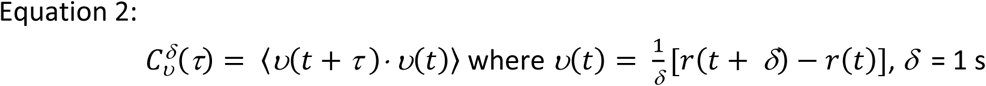

For confinement radius analysis, the same trajectory length thresholds were used. First, the average position of the track was calculated by averaging the xy coordinates. The average of the difference between this position and the track coordinates gives the confinement radius.

#### Carboxysome localization heatmaps

A custom Jupyter notebook was used to perform this analysis: *Spideymaps_carboxysomes_multi.ipynb*. The localization coordinates of detected carboxysome foci and cell phase masks were used to make localization density heatmaps with the *spideymaps* tool in Python. First, the *colicoords* (Smit et al., 2019) package was used to convert the Cartesian coordinates of each cell into cellular coordinates that better describe the rod-shaped cell geometry. Then, cells were rotated and scaled to generate a uniform coordinate plane for all cells in a population. Localizations were binned and symmetrized across the *x* and *y* axes. Localization counts and associated bin areas were summed across all cells, and the summed number of counts was divided by the summed area to calculate the localization density (counts per pixel^2^). The Spideymaps package can be found in the Biteen Lab Github repository: https://github.com/BiteenMatlab/spideymaps.

#### Kymographs

Time-lapse fluorescence microscopy image stacks of phase contrast (200-ms exposure) and mNG (2-s exposure) were collected at 10-s intervals and drift corrected as above. The corrected phase contrast image was opened in Fiji, and a line was manually drawn along the long axis of a cell. The multi-kymograph tool was used to extract the intensity profile with a 9-pixel (594 nm) width. The resulting kymograph was saved as a TIFF file. For visualization, *norm_kymographs.py* script was used. The intensity profile at each timepoint of the kymograph is normalized to the minimum and maximum intensity values within the corresponding time point, thus the values are normalized to 0 and 1. This post-processing removed the effects of photobleaching in the visualization of McdA-mNG.

#### Nucleoid morphology

The cell phase masks were used to determine the pixel intensities within the inner rim of the cell boundary (*skimage.segmentation.find_boundaries*). The distribution of the single cell inner rim pixel intensities was fit to a Gaussian distribution (*lmfit.Model.fit*) to determine the average intensity of the cytoplasmic region of the cell. This intensity was subtracted across all pixels in the cell. A Gaussian blur (SD of Gaussian = 1 pixel (66 nm); *skimage.filters.gaussian*) was applied to the background-subtracted image. Next, the nucleoid region was detected using Yen’s thresholding (*skimage.filters.threshold_yen*) (Yen et al., 1995). The length of the nucleoid is the length of the long axis of the segmented region.

### Single-molecule fluorescence microscopy

Cells expressing native protein fusions to PAmCherry were inoculated into S-6 medium and grown with shaking (130 rpm) at 30 °C, and in air supplemented with 5% CO_2_ for 30 – 36 h before back diluting 1:100-200 in 5 mL of fresh S-6 medium. Cells were then grown under the same conditions for 18 – 24 h. Prior to imaging, 1 – 1.5 mL of cells were harvested by centrifugation at 5000 ×g for 5 min and resuspended in 10 – 20 µL of fresh S-6 medium. Agarose pads were made at 2% (w/v) with S-6 medium without thiosulfate. An aliquot of 2 µL of cells was loaded onto an agarose pad and sandwiched between two coverslips. Samples were allowed to equilibrate on the stage top for 10 – 15 min prior to image collection. Cells were imaged at room temperature with a custom-built microscope equipped with a 100×, 1.40 numerical aperture oil-immersion objective. For fast single-molecule tracking experiments, 100 – 200 ms doses of 405-nm laser (Coherent Cube 405-100; 1.5 W/cm^2^) were used for PAmCherry photoactivation, and a 561-nm laser (Coherent-Sapphire 561-50; 0.6 kW/cm^2^) was used for imaging. The fluorescence emission was passed through a tri-band dichroic mirror and emission filter, and 4000 – 6000 frames were collected at a rate of 50 Hz using a 512 × 512-pixel Photometrics Evolve EMCCD camera. Samples were imaged for 1 – 1.5 hr before loading a new sample.

### Single-molecule data analysis

Phase-contrast microscopy images were used to segment bacterial cells (see Image analysis for details) prior to single-molecule localization and tracking. Image stacks with drift were excluded from analysis. Single molecules were detected and localized using the SMALL-LABS package in MATLAB (Isaacoff et al., 2019). Photoactivation by 406 nm light excites CbbS-mTQ, therefore an intensity-based threshold was set on the mean intensity of each frame to exclude the photoactivation frames from localization analysis. Putative single molecules were identified as non-overlapping punctate spots if an area of 8 × 8 pixels had pixel intensities above the 98^th^ percentile intensity of the frame. Localization was done by fitting the putative single molecule spot to a 2D Gaussian; ‘good’ localization conditions were defined by a spot width = 7 pixels (343 nm), a tolerance threshold of 2.5 on the SD of the 2D Gaussian fit, and fit error ≤ 0.065 (Isaacoff et al., 2019). Good localizations were then assembled into trajectories via the Hungarian algorithm (Munkres, 1957) using a maximum step size of 0.980 µm. Trajectories of at least 5 frames in which the molecule was localized in more than 50% of the frames and for which there were no more than 3 consecutive dark frames (in which no molecule was localized) were allowed.

Each trajectory was analyzed independently to extract a trajectory-average apparent diffusion coefficient, *D_app_*, and the associated localization uncertainty, *σ*. The mean-squared displacement, <*r^2^*>, as a function of time lag (*nτ*), where *n* is an integer and *τ* is the 20 time between frames, was calculated for up to the first 3 *τ* (20 – 60 ms) and fit with a weighted least-squares fit (*scipy.optimize.curve_fit*) to Equation 3. The slope of this curve is proportional to *D_app_* according to a modified version of the diffusion model in two dimensions (Equation 3). The modified diffusion model was used to account for motion blur due to averaging the true position of a molecule over the time of a single acquisition frame (Berglund, 2010). The following bounds were placed on the fit parameters: *D_app_* [0, infinity]; *σ* [0, infinity]. Trajectories with an *R*^2^ ≤ 0.2 threshold were excluded from further analysis. The distribution *D_app_* was then fit to a GMM (*sklearn.mixture.GaussianMixture*) with two components, which provides the average *D_app_*, the standard deviation, and the population weight associated with each component. Bootstrap analysis was performed by randomly sampling the dataset with repetition to generate a set with a sample size equal to that of the original. Reported means and standard deviations are from independent experiments (Table S1).

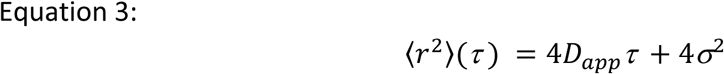

### Estimation of localization uncertainty

To estimate the localization precision of immobile PAmCherry molecules in *H. neapolitanus* under the imaging conditions used, we performed the same single-molecule experiments with fixed cells. *H. neapolitanus* cells containing PAmCherry-McdB molecules expressed off an inducible promoter were induced with 0.5 mM IPTG for 3 h. Fixation was achieved by harvesting 1.5 mL of culture, concentrating the cells to ∼ 20 µL, and adding formaldehyde: PBS to a final concentration of 2.5 % (v/v). The sample was then incubated at room temperature for 10 min before storage at 4 °C for 18 h. Imaging, localization, and tracking were done using the conditions described for 20-ms and 80-ms imaging. The confinement radii for tracks with at least five frames were calculated. The radius distribution was fit to a Gaussian function (Equation 4) (*lmfit.Model.fit*) where *r* is the confinement radius, *σ* is the localization uncertainty, and *ω* is the standard deviation associated with *σ*. Analysis resulted in an uncertainty of 29 ± 6 nm for 20 ms imaging and 19 ± 6 nm for time-lapse (Supplementary Fig. 9).

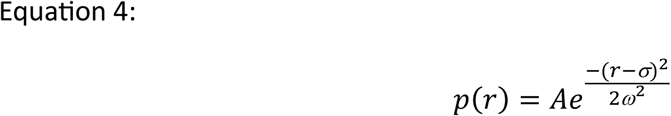

### Single-molecule time-lapse fluorescence microscopy

Single-molecule time-lapse experiments were performed as described above, but with modifications. Camera exposure times were increased to 80 ms to bias the detection to slow McdA-PAmCherry^SW^ molecules (i.e., the subset of molecules bound to DNA). Imaging was done with continuous acquisition or with an added dark-time delay between frames (140, 340, 640, or 840 ms). The 561-nm excitation power was lowered to 0.2 kW/cm^2^ to reduce the rate of photobleaching. Localization and tracking of PAmCherry constructs were performed with SMALL-LABS. SMALL-LABS parameters were modified to identify single molecules more strictly and prevent localizations of blurred molecules. The tolerance threshold was decreased to 1.75 on the SD of the 2D Gaussian fit, and the error on the fit was changed to ≤ 0.125. For the continuous imaging dataset, the max localization linking distance was set to 8 pixels (392 nm), and a maximum of 1 dark frame gap was allowed. The localization linking distance for each time-lapse experiment was determined by calculating the maximum possible squared displacement given the measured diffusion coefficient of the mobile state from the continuous imaging dataset: 0.018 ± 0.004 µm^2^s^−1^ and the time-lapse, *τ*, between frames. A one-pixel (49-nm) buffer was added to account for noise.

Imaging of ParB-mNG foci was performed on the same microscope setup as PAmCherry imaging. Here, all ParB-mNG molecules were excited simultaneously, as there is no photoactivation mechanism. To maintain a comparable localization uncertainty, ParB-mNG molecules were excited with 0.01 W/cm^2^ of 488-nm light. A total of 50 frames was collected using 80-ms camera integration times and a 140-ms dark-time delay. Localization and tracking were performed with TrackMate. Fluorescent spots were detected with LoG (radius = 6 pixels or 294 nm) using subpixel localization. Localizations were then linked with the LAP tracker method using a maximum linking length of 5 pixels (245 nm) and no gaps allowed since the focus signal was stable throughout the imaging period.

Confinement radius analysis was performed as above using the entire trajectory length for tracks with at least 4 localizations.

### Cumulative distribution function (CDF) analysis

Trajectories were filtered to a minimum length of 4 localizations and at most 1 gap between frames. The squared displacement, *r^2^*, was calculated for consecutive localizations within a trajectory (*τ* = 220 ms). The cumulative probability of the squared displacements in the observation period *P*(*r^2^, τ*) was generated from the pool of squared displacements across multiple tracks for each construct replicate by counting the number of squared displacements less than or equal to *r^2^* normalized by the sample size. The CDF of *r^2^* was fit to analytical functions (Equations 5 and 6) (*lmfit.Model.fit*) describing the diffusive processes with one or two dynamic states. *D_1_* and *D_2_* are the diffusion coefficients for the different states, and *α* describes the relative fraction between the states. Here, the *D_1_*term is assumed to be immobile and was constrained to a max value of 0.0015 µm^2^s^−1^, the theoretical measurable diffusion coefficient lower bound based on the uncertainty. The minimum for the *D_2_* term (mobile) was constrained to the same value. The error term is 4*σ^2^*, with *σ* = 0.019 µm, which was obtained as described above (Supplementary Fig. 9). Based on the calculated residuals, the two-state model resulted in the best fit (Supplementary Fig. 10).

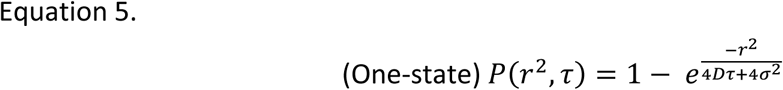

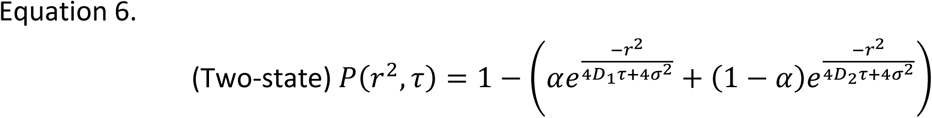

### Single-molecule residence time *in vivo*

The residence time of McdA binding to DNA was attained by analyzing the single-molecule time-lapse datasets as previously described (Chen et al., 2024) based on modeling the binding as a direct two-component association/dissociation reaction: *AB* ⇋ *A* + *B*. The measured residence time of each McdA-PAmCherry^SW^ molecule is estimated from the lifetime of the tracked fluorescence signal. *k_app_*is obtained by fitting the survival curve (*lmfit.Model.fit*), *P* (Equation 7), of the measured residence times, *τ_measured_*, to a single exponential decay function:

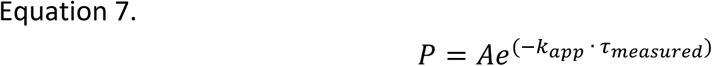

Under the experimental conditions, the measured apparent dissociation rate, *k_app_*, contains contributions from the true dissociation rate, *k_diss_*, and the photobleaching rate of the PAmCherry label, *k_bleaching_*. The *k_bleaching_* was measured by collecting data at multiple delay times, *τ_delay_*, while maintaining a constant camera integration time, *τ_int_*, to keep *k_bleaching_*constant for the different time delays, *τ_TL_ = τ_int_ + τ_delay_*. The residence time, *τ_measured_ = nτ_TL_*, was determined by measuring the length, *n*, of each trajectory with 1 gap allowed. The true dissociation rate, *k_diss_*, was estimated from a linear regression (*scipy.optimize.curve_fit*) of the two-term relationship:

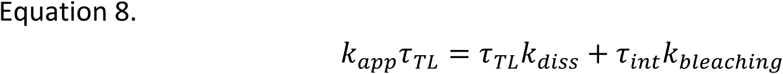

The linear regression was weighted by the uncertainty of each data point. Uncertainty values of each point are the error on the exponential fit for each time-lapse. The error in the linear regression is the uncertainty in *k_diss_*.

### Quantification of McdA copy number

#### Measurement of single mNG fluorescence

To measure the intensity of single mNG molecules in *H. neapolitanus*, cells expressing mNG-FliN were grown and prepared for imaging as in the “Single-molecule fluorescence microscopy” section. Here, the approach was to measure single mNG photobleaching events since mNG-FliN forms a single fluorescent focus at the cell pole. Cells were illuminated with a 488-nm laser (Coherent Cube 488-50; 4.5 W/cm²) and a continuous series of frames (50 or 100) was captured using 2-s integration times. To quantify the intensity of single mNG molecules, first, every imaging frame was corrected for the Gaussian profile of the 488-nm beam to account for differences in excitation. Next, fluorescent spots were detected and their integrated intensity counts collected over the entire imaging period. To determine the change in the spot intensity as a result of a single photobleaching event, a semi-automated approach was taken. Single-spot fluorescence intensity traces (*N* = 88 traces) were manually annotated to note when the photobleaching events occurred; then the average of the seven frames pre- and post-the photobleaching frame was used to calculate the change in the spot intensity as a result of a single mNG photobleaching. The distribution of these intensities (*N* = 192 photobleaching events) was then fit to a Gaussian model and resulted in 11,245 ± 4094 intensity counts per single mNG molecule.

#### Measurement of total mNG-McdA fluorescence per cell

*H. neapolitanus* cells expressing CbbS-mTQ or mNG-McdA^SW^ and CbbS-mTQ were grown and prepared for imaging as in the section “Single-molecule fluorescence microscopy”. Cells were illuminated with a 488-nm laser (Coherent Cube 488-50; 4.5 W/cm²) and single frames were captured using a 2-s integration time. Phase contrast images were captured for subsequent cell segmentation done as described in the “Image processing and analysis” section. Fluorescence images were corrected for the Gaussian shape of the 488-nm beam. Using the cell segmentations, the cell pixel fluorescence intensities were summed to determine the total cellular intensity. To correct for the cell background (autofluorescence and CbbS-mTQ bleed through), the average cellular intensity of CbbS-mTQ cells was calculated and used as background. The number of mNG-McdA^SW^ molecules per cell was calculated as the ratio between the background-corrected mNG-McdA^SW^ fluorescence per cell and the mean fluorescence of single mNG-FliN molecules.

### DNA-McdA-McdB-carboxysome complex dissociation constant estimation

To calculate the dissociation rate of the McdA-McdB bonds, *k_off_,* first the rate of association of McdA to the nucleoid was estimated using the experimentally measured equilibrium dissociation constant, *K_D_*, and the dissociation rate of McdA from the nucleoid, *k_diss_*. Here *k_on_* = *k_off_* / *K_D_* = 3.71 x 10^4^ M^−1^s^−1^. Assuming that *k_on_* of McdA remains unchanged in the presence of McdB (Kohler et al., 2022), the *k_off_* of McdA from the nucleoid as a result of its interaction with McdB can be estimated by *k_off_* = *K_D_k_on_* = 0.328 s^−1^.

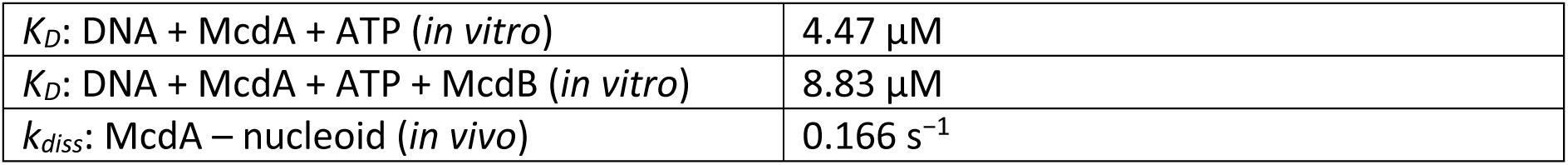

|  |  |
| --- | --- |
| $K_D$: DNA + McdA + ATP (*in vitro*) | 4.47 $\mu\text{M}$ |
| $K_D$: DNA + McdA + ATP + McdB (*in vitro*) | 8.83 $\mu\text{M}$ |
| $k_{\text{diss}}$: McdA – nucleoid (*in vivo*) | 0.166 $\text{s}^{-1}$ |

### The mechanochemical Brownian ratchet modeling

The details of the Brownian ratchet force-generation mechanism, as well as the implementation of stochastic simulation of the reaction-diffusion processes involved in cargo movement, can be found in our previous publications (Hu et al., 2015; Hu et al., 2017a; Hu et al., 2017b; MacCready et al., 2018; Hu et al., 2021; Byrne et al., 2025). In this study, wherever possible, we performed the simulations by incorporating the experimentally measured parameters of the *H. neapolitanus* McdAB system (Table S4). We estimated the number of McdA molecules per cell (Supplementary Fig. 11) and the dimensions of the nucleoid (Supplementary Fig. 12) using quantitative fluorescence microscopy. Additionally, we estimated the rate of dissociation of the DNA-McdA-McdB-carboxysome complex using the equilibrium DNA-dissociation constants in the presence and absence of McdB (Figure 4c) and the (5) *in vivo* McdB-independent dissociation rate of McdA from the nucleoid (Figure 4f).

For each simulated system with a fixed set of model parameters (e.g., carboxysome number), we ran the simulation for 300 seconds to generate individual trajectories for each carboxysome in the system. We then calculated the extent of equal partitioning and the average segregation distance based on the carboxysome positions at the end of the simulation. The extent of equal partition is defined as the weighted probability min(*n_l_*, *n_r_*) ⁄ max(*n_l_*, *n_r_*), where *n_l_* is the number of carboxysomes on the left half of the nucleoid and *n_r_* is the number of carboxysomes located on the right of the nucleoid at the end of the simulation. The coefficient of variation of the extent of equal partitioning is defined as the ratio of the standard deviation and mean of the extent of equal partitioning.

## ACKNOWLEDGEMENTS

We are grateful to Jose M. Limcaoco for running the McdA-PAmCherry^SW^ sequence through AF2 and Cade T. Harkner for preliminary time-lapse fluorescence imaging experiments. We thank Dr. Luis Ortíz-Rodriguez for time-lapse fluorescence imaging support and Dr. Daniel Foust for help with Spideymaps analysis. C.A.A. was funded by the University of Michigan Rackham Predoctoral Fellowship. A.G.V. acknowledges funding from the National Institute of General Medical Sciences of the National Institutes of Health (grant R35-GM152128), and J.S.B. acknowledges funding from the National Institute of General Medical Sciences of the National Institutes of Health (grant R01-GM144731).

## AUTHOR CONTRIBUTIONS

Conceptualization: C.A.A., A.G.V, and J.S.B., Methodology: C.A.A., H.M.S, L.H., and L.T.P., Software: C.A.A., Formal analysis: C.A.A., H.M.S., and L.H., Investigation: C.A.A., H.M.S., L.H., and H.R., Resources: J.L., A.G.V and J.S.B., Data curation: C.A.A. Writing – original draft: C.A.A., A.G.V., J.S.B., Writing – review and editing: C.A.A., A.G.V., and J.S.B., Supervision: J.L, A.G.V and J.S.B.

