## Supplemental Information for "The kinetics and mobility of a ParA ATPase drive carboxysome distribution in *Halothiobacillus neapolitanus*"

### Contents

- **Supplementary Tables S1 – S9**
- **Supplementary Figures S1 – S12**

### SUPPLEMENTARY TABLES

**Table S1.** Apparent diffusion coefficients, $D_{app}$, from two-state Gaussian Mixture fitting of data shown in Figure 4d-e and Supplementary Fig. 8.

| Sample | *N* tracks | Slow (%) | $D_{app}$ (µm²/s) | Fast (%) | $D_{app}$ (µm²/s) |
|---|---|---|---|---|---|
| WT (rep 1) | 4524 | 45.9 | 0.0115 | 55.1 | 0.3729 |
| WT (rep 2) | 3481 | 55.8 | 0.0113 | 44.2 | 0.3993 |
| WT (rep 3) | 4221 | 61.6 | 0.0102 | 38.4 | 0.3728 |
| WT (rep 4) | 3665 | 56.1 | 0.0129 | 43.9 | 0.4035 |
| Δ*mcdB* (rep 1) | 4165 | 60.6 | 0.0081 | 39.4 | 0.1630 |
| Δ*mcdB* (rep 2) | 3829 | 65.5 | 0.0110 | 34.5 | 0.2542 |
| Δ*mcdB* (rep 3) | 2652 | 63.2 | 0.0080 | 36.8 | 0.2110 |
| Δ*mcdB* (rep 4) | 4853 | 63.1 | 0.0106 | 36.9 | 0.3362 |
| Δ*mcdB* (rep 5) | 4598 | 77.0 | 0.0110 | 23.0 | 0.2744 |
| McdA[K20A] (rep 1) | 4403 | 16.5 | 0.0236 | 83.5 | 0.3781 |
| McdA[K20A] (rep 2) | 2915 | 12.1 | 0.0228 | 87.9 | 0.4234 |
| McdA[K20A] (rep 3) | 4440 | 16.9 | 0.0277 | 83.1 | 0.4317 |
| McdB[W94G] (rep 1) | 4364 | 65.7 | 0.0088 | 34.3 | 0.2626 |
| McdB[W94G] (rep 2) | 5527 | 67.5 | 0.0110 | 32.5 | 0.2384 |

**Table S2.** Confinement radii for data shown in Figure 5b.

| Sample | *N* tracks | Confinement radius (nm) |
|---|---|---|
| ParB (rep 1) | 225 | 29.5 |
| ParB (rep 2) | 325 | 28.6 |
| ParB (rep 3) | 400 | 27.4 |
| WT (rep 1) | 898 | 46.6 |
| WT (rep 2) | 562 | 51.9 |
| WT (rep 3) | 907 | 42.6 |
| Δ*mcdB* (rep 1) | 956 | 51.2 |
| Δ*mcdB* (rep 2) | 470 | 62.4 |
| Δ*mcdB* (rep 3) | 564 | 59.5 |

**Table S3.** Results from two-state CPD fitting for data shown in Figure 5c-d and Supplementary Fig. 10.

| Sample | Time lapse (ms) | *N* tracks (displacements) | Immobile (%) | $D_{app}$ (µm²/s) | Mobile (%) | $D_{app}$ (µm²/s) |
|---|---|---|---|---|---|---|
| Δ*mcdB* (rep 1) | 80 | 1606 (12641) | 62.8 | $1.2 \times 10^{-3}$ | 37.2 | $1.6 \times 10^{-2}$ |
| Δ*mcdB* (rep 2) | 80 | 1496 (13078) | 50.1 | $1.0 \times 10^{-4}$ | 49.9 | $1.0 \times 10^{-2}$ |
| Δ*mcdB* (rep 3) | 80 | 1385 (11818) | 50.0 | $1.1 \times 10^{-4}$ | 50.0 | $1.0 \times 10^{-2}$ |
| WT (rep 1) | 220 | 898 (8768) | 61.8 | $1.0 \times 10^{-4}$ | 38.2 | $6.2 \times 10^{-3}$ |
| WT (rep 2) | 220 | 562 (5198) | 52.2 | $1.0 \times 10^{-4}$ | 47.8 | $5.9 \times 10^{-3}$ |
| WT (rep 3) | 220 | 907 (6192) | 54.8 | $1.0 \times 10^{-4}$ | 45.2 | $5.0 \times 10^{-3}$ |
| Δ*mcdB* (rep 1) | 220 | 956 (7493) | 35.6 | $1.0 \times 10^{-4}$ | 64.4 | $6.0 \times 10^{-3}$ |
| Δ*mcdB* (rep 2) | 220 | 470 (4037) | 39.6 | $5.8 \times 10^{-4}$ | 60.4 | $7.9 \times 10^{-2}$ |
| Δ*mcdB* (rep 3) | 220 | 564 (3665) | 30.2 | $1.0 \times 10^{-4}$ | 69.8 | $7.4 \times 10^{-2}$ |
| ParB-mNG (rep 1) | 220 | 225 (10465) | 30.4 | $2.7 \times 10^{-4}$ | 69.6 | $1.5 \times 10^{-3}$ |
| ParB-mNG (rep 2) | 220 | 325 (15791) | 51.9 | $3.1 \times 10^{-4}$ | 48.1 | $1.7 \times 10^{-3}$ |
| ParB-mNG (rep 3) | 220 | 400 (19506) | 53.2 | $2.7 \times 10^{-4}$ | 46.8 | $1.4 \times 10^{-3}$ |

**Table S4.** Experimentally measured parameters of the *H. neapolitanus* McdAB system.

| Parameter | Value range | Reference |
|---|---|---|
| Rate of McdA-McdB bond formation | 3300.0 $s^{-1}$ | Hu et al., 2021 |
| Rate of McdA-McdB bond dissociation | 0.33 $s^{-1}$ | This study |
| Rate of cytosolic McdA·ATP binding to nucleoid | 3.0 $s^{-1}$ | Hu et al., 2021 |
| Rate of McdA·ATP dissociation from the nucleoid | 0.01 $s^{-1}$ | Hu et al., 2021 |
| Rate of McdA·ADP dissociation from the nucleoid | 5.0 $s^{-1}$ | Hu et al., 2021 |
| Diffusion constant of McdA·ATP along the nucleoid | 0.01 $\mu m^2 s^{-1}$ | This study |
| Number of McdA Dimers per cell | 0 – 2000 | This study |
| Length of the nucleoid | 0.96 µm | This study |
| Width of the nucleoid | 0.62 µm | This study |
| Diameter of carboxysomes | 150 nm | Iancu et al., 2010 |
| Carboxysome number | 2 – 12 | Iancu et al., 2010 |

**Table S5.** Strains used in this study.

| Strain | Genotype | Source |
|---|---|---|
| AGV_Hn2 | *Hn::cbbS-mTQ* | MacCready and Tran et al., 2021 |
| AGV_Hn8 | *Hn::mNG-fliN* | Pulianmackal et al., 2023 |
| AGV_Hn14 | *Hn::parB-mNG* | Pulianmackal et al., 2023 |
| AGV_Hn100 | *Hn::cbbS-mTQ:: ΔmcdA* | MacCready and Tran et al., 2021 |
| AGV_Hn321 | *Hn::cbbS-mTQ:: pTrc-PAmCherry-mcdB* | This study |
| AGV_Hn431 | *Hn::cbbS-mTQ:: mcdA-PAmCherry*$^{SW}$ | This study |
| AGV_Hn432 | *Hn::cbbS-mTQ:: ΔmcdAB* | This study |
| AGV_Hn441 | *Hn::cbbS-mTQ:: mcdA-PamC*$^{SW}$ *+ ΔmcdB* | This study |
| AGV_Hn481 | *Hn::cbbS-mTQ:: mcdA[K20A]-mNG*$^{SW}$ | This study |
| AGV_Hn498 | *Hn::cbbS-mTQ:: mcdA-mNG*$^{SW}$ *+ ΔmcdB* | This study |
| AGV_Hn404 | *Hn::cbbS-mTQ:: PamC-mcdA* | This study |
| AGV_Hn518 | *Hn::cbbS-mTQ:: mcdA-PamC* | This study |
| AGV_Hn532 | *Hn::cbbS-mTQ:: mcdA-mNG*$^{SW}$ | This study |
| AGV_Hn537 | *Hn::CbbS-mTQ:: mcdA[K20A]-PamC*$^{SW}$ | This study |
| AGV_Hn554 | *Hn::CbbS-mTQ:: mcdA-PamC*$^{SW}$ *+ mcdB[W94G]* | This study |

**Table S6.** Plasmids used in this study

| Plasmid name | Relevant genetic elements | Source |
|---|---|---|
| For constructing *H. neapolitanus* strains | | |
| pLT04 | *Hn0910-0911_specR_Hn0913-0914* | MacCready and Tran et al., 2021 |
| pLT17 | *pTrc-mNeonGreen-mcdB-kanR* | MacCready and Tran et al., 2021 |
| pLT67 | *pTrc-PAmCherry-mcdB-kanR* | This study |
| pLT79 | *native-PAmCherry-mcdA-mcdB-chlR* | This study |
| pLT87 | *native-mcdA-PAmCherry-mcdB-chlR* | This study |
| pLT90 | *Hn0910_specR_Hn0913-0914* | This study |
| pLT94 | *native-mcdA*$^{1-59}$*-PAmCherry-mcdA*$^{60-216}$*-mcdB-chlR* | This study |
| pLT103 | *native-mcdA*$^{1-59}$*-PAmCherry-mcdA*$^{60-216}$*-chlR* | This study |
| pLT104 | *native-mcdA*$^{1-59}$*[K20A]-PAmCherry-mcdA*$^{60-216}$*-mcdB-chlR* | This study |
| pCA2 | *native-mcdA*$^{1-59}$*-mNeonGreen-mcdA*$^{60-216}$*-mcdB-chlR* | This study |
| pCA3 | *native-mcdA*$^{1-59}$*-mNeonGreen-mcdA*$^{60-216}$*-chlR* | This study |
| pCA4 | *native-mcdA*$^{1-59}$*[K20A]-mNeonGreen-mcdA*$^{60-216}$*-mcdB-chlR* | This study |
| pCA31 | *native-mcdA*$^{1-59}$*-PAmCherry-mcdA*$^{60-216}$*-mcdB[W94G]-chlR* | This study |
| For transformation into *E. coli* BL21 (DE3) | | |
| pJB26 | pET11b::*hisx6-sumo-mcdB-ampR* | Basalla et al., 2023 |
| pAV96 | pET15b::*hisx6-Hn_0912-GFP-ampR* | This study |
| pHS02 | pET15b::*hisx6-SUMO-mcdA-ampR* | This study |
| pHS06 | pET15b::*hisx6-SUMO-mcdA[K20A]-ampR* | This study |
| pHS08 | pET15b::*hisx6-SUMO-mcdA[R158E]-ampR* | This study |
| For EMSA | | |
| pUC19 | | New England Biolabs (Cat. # N3041S) |

**Table S7.** DNA oligonucleotides used in this study.

| Oligo name | Sequence (5’ to 3’) | Purpose |
|---|---|---|
| ptrc_backbone_rev | ATGTTTTTCCTCCTTGTGTGAAATTGTTAT | To make pLT67 |
| mcdB_CO_fwd | ATGACGAATCTGGAAGACAAA | To make pLT67 |
| PAmCherry_fwd | GTGAGCAAGGGCGAGGAG | To make pLT67, pLT79 |

| | | |
|---|---|---|
| PAmCherry_rev | CTTGTACAGCTCGTCCATGC | To make pLT67 |
| Hn_0913_rev | TGTTATCCTCCTCGCCCTTGCTCACGGCGTTAGCTATACTTGGCG | To make pLT79 |
| mcdA/mcdB_CO_fwd | TCTTAAATGAAGTCATCCCATCATGACGAATCTGGAAGACAAATTATCA | To make pLT79 |
| mcdA_CO_rev | CATGATGGGATGACTTCATTTAAGATC | To make pLT79 |
| GSGx9-PAmCherry_fwd | ATCAGGTGGCGGTGGCGGTGGTGGCGGTGGTGTGAGCAAGGGCGAGGAG | To make pLT87 |
| pamcherry_rev_2 | TTTGTACAGCTCGTCCATGCC | To make pLT87 |
| PAmCherry_mcdB | CCGGCGGCATGGACGAGCTGTACAAATGACTAATTTAGAAGATAAACTGAGTG | To make pLT87 |
| GSGx9_mcdA_rev | GCCACCACCGCCACCGCCACCTGATCCTGACGGGATAACCTCGTTAAG | To make pLT87 |
| Pterin_rev | TTAGCTATACTTGGCGGCCAAG | To make pLT90 |
| Hn0910_fwd | GTCAAACTATGAACCCAGCGAC | To make pLT90 |
| GS/mcdA_fwd | GTACAAGGGTTCTGGTAGTGGATCTGACGCTCAAGTCCCCGTC | To make pLT94 |
| Hn0912_GS_rev | TGCTCACTGATCCTGATCCTGAACCCGCCTCTTTGCTTCCCCA | To make pLT94, pCA2, pCA3, pCA4 |
| GSGSGS_PAmCherry_fwd | GGTTCAGGATCAGGATCAGTGAGCAAGGGCGAGGAG | To make pLT94 |
| GSGSGS_PAmcherry_rev | AGATCCACTACCAGAACCCTTGTACAGCTCGTCCATGC | To make pLT94 |
| delmcdB_fwd | GTTATCCCGTCATGATTTTGTCCGAACATCGCGAATTACG | To make pLT103 |
| delmcdB_rev | TCATGACGGGATAACCTCGTTAAGG | To make pLT103 |
| SDM_McdA_K20A_fwd_long | AAAGGTGGCTGTGGCGCCACGACC | To make pLT104, pCA4 |
| mcdA_K20R_rev_long | GCCACAGCCACCTTTGAGGTTGGCGACA | To make pLT104, pCA4 |
| GS-mNG_fwd | GGTTCAGGATCAGGATCAATGGTGTCGAAAGGAGAAGAAG | To make pCA2, pCA3 |
| mNG-GS_rev | AGATCCACTACCAGAACCCTTGTATAATTCGTCCATCCCC | To make pCA2, pCA3 |
| GS_mcdA_fwd_2 | GGTTCTGGTAGTGGATCTGACGCTCAAGTCCCCGT | To make pCA2, pCA3 |
| SDM_McdB_W94G_fwd | TCGCCGTGTAGGCCCAGATTGAT | To make pCA31 |
| SDM_McdB_W94G_rev | GGATGCGTCGATGTG | To make pCA31 |

| | | |
|---|---|---|
| 1_His SUMO Forward | CTTTAAGAAGGAGATATACATATGGGCAGTAGCCACCATC | To make pHS02 |
| 2_SUMO w HN-McdA tail Reverse | GCTTTATTGCTGGCCATGCCCCCGATCTGTTCACGGTGTG | To make pHS02 |
| 3_SUMO tail w HN McdB backbone Forward | GTGAACAGATCGGGGGCATGGCCAGCAATAAAGCATTTAC | To make pHS02 |
| 4_RBS His pET15b backbone Reverse | CTACTGCCCATATGTATATCTCCTTCTTAAAGTTAAAC | To make pHS02 |
| 30_Hn-McdA K20A Forward | TGGTTGTGGTgcgACCACCATTAG | To make pHS06 |
| 31_Hn-McdA K20A Reverse | CCTTTCAGATTTGCAACTG | To make pHS06 |
| 36_Hn-McdA R158E Forward | CGCAATGACCgaaGCAATGCAGAC | To make pHS08 |
| 37_Hn-McdA R158E Reverse | CTACGCGGTTCGGTCTGATTCAG | To make pHS08 |

**Table S8.** Chemicals used in this study.

| Chemical | Source | Catalog number |
|---|---|---|
| 4',6-diamidino-2-phenylindole (DAPI), dihydrochloride | ThermoFisher Scientific | D1306 |
| Isopropylthio-β-galactoside (IPTG) | ThermoFisher Scientific | 15529019 |
| EnzChek Phosphatase Assay Kit | ThermoFisher Scientific | E12020 |

**Table S9.** Software used in this study.

| Program / library / script | Package | Source |
|---|---|---|
| Fiji | imagej.net/software/fiji | Scheneider et al., 2012 |
| Trackmate | imagej.net/plugins/trackmate | Tinevez et al., 2017 |
| MATLAB (2023b) | mathworks.com/products/new_products/release2023b.html | Mathworks |
| ColabFold | colab.research.google.com/github/sokrypton/ColabFold/blob/main/AlphaFold2.ipynb | Mirdita et al , 2022 |
| PyMOL | pymol.org (v 3.0.1) | Schrodinger, Inc. |
| Python 3 | python.org (v 3.12.5) | Python Software Foundation |
| Scikit-image | scikit-image.org | Van der Walt et al., 2014 |

| | | |
|---|---|---|
| Scikit-learn | scikit-learn.org/stable | Pedregosa et al., 2012 |
| Matplotlib | matplotlib.org | Hunter, 2007 |
| Scipy | scipy.org | Virtanen et al., 2020 |
| Seaborn | seaborn.pydata.org | Waskom, 2021 |
| Pandas | pandas.pydata.org | McKinney et al., 2010 |
| LMFIT | lmfit.github.io/lmfit-py | Newville et al., 2014 |
| Omnipose | github.com/kevinjohncutler/omnipose | Cutler et al., 2022 |
| Spideymaps | github.com/BiteenMatlab/spideymaps | Dr. Daniel Foust |
| PhaseMasks_omni.py | Cell segmentation by Omnipose; script for batch processing | This study |
| cell_registration.py | To drift correct image stacks. Requires specific package versions: scikit-image 0.16.2, numpy 1.18.5 | This study |
| detect_carboxysomes.py | To localize carboxysome foci, determine their nearest neighbor distance, and measure the associated cellular morphology | This study |
| Spideymaps_carboxysomes_multi.ipynb | To make the carboxysome localization heatmaps | This study |
| norm_kymographs.py | To normalize kymographs and generate plot | This study |
| nucleoid_morphology.py | To segment nucleoid regions and measure the associated cellular morphology | This study |
| SMALLLABS_main_acti.m | To localize and track single molecules with activation frames excluded | This study |
| get_ATPturnover_rates.py | To calculate ATP turnover rate | This study |
| get_specific_activity.py | To calculate ATPase specific activity | This study |
| copy_calculate_protein_copy.py | To calculate single-cell protein copy numbers and concentration using fluorescence imaging data | This study |
| copy_collect_traces.py | To detect fluorescent spots in time series data and retrieve intensity traces | This study |
| copy_trace_analysis.py | To plot intensity traces for semi-automated analysis of photobleaching events | This study |
| copy_norm_stacks.py | To normalize fluorescence images with a corresponding laser beam profile | This study |
| copy_singlecell_intensity.py | To calculate single-cell fluorescence intensities | This study |
| calc_MSD_2D_for_SMALLL | To calculate the diffusion coefficient by | This study |

| | | |
|---|---|---|
| ABS.py | MSD analysis using SMALL-LABS detected tracks | |
| calc_MSD_2D_for_trackmate.py | To calculate the diffusion coefficient by MSD analysis using TrackMate detected tracks | This study |
| calc_VACF_2D.py | To calculate the velocity autocorrelation function of a trajectory | This study |
| calc_localization_uncertainty.py | To calculate the localization uncertainty associated with a fixed-sample trajectory | This study |
| calc_radius_confinement_carboxysomes.py | To calculate the radius of confinement of carboxysome trajectories | This study |
| calc_radius_confinement_singlemols.py | To calculate the radius of confinement of single-molecule trajectories | This study |
| calc_track_CPD_smalllabs.py | To calculate the diffusion coefficient by CPD analysis using SMALL-LABS detected tracks | This study |
| calc_track_CPD_trackmate.py | To calculate the diffusion coefficient by CPD analysis using TrackMate detected tracks | This study |
| calc_track_lifetimes.py | To calculate the ‘On’ times of single-molecule trajectories to determine residence times | This study |

### SUPPLEMENTARY FIGURES

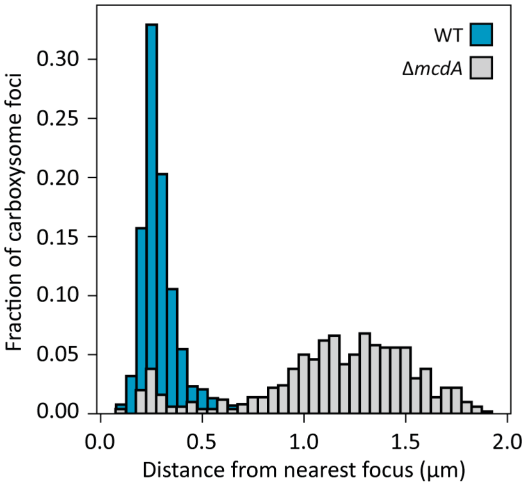


**Supplementary Fig. 1: Distribution of distances between adjacent carboxysome foci**. Distribution of the distances between each carboxysome focus and its nearest neighbor in WT (*N* = 3477 foci) and Δ*mcdA* (*N* = 500 foci) cells. Only cells with >1 focus were analyzed.

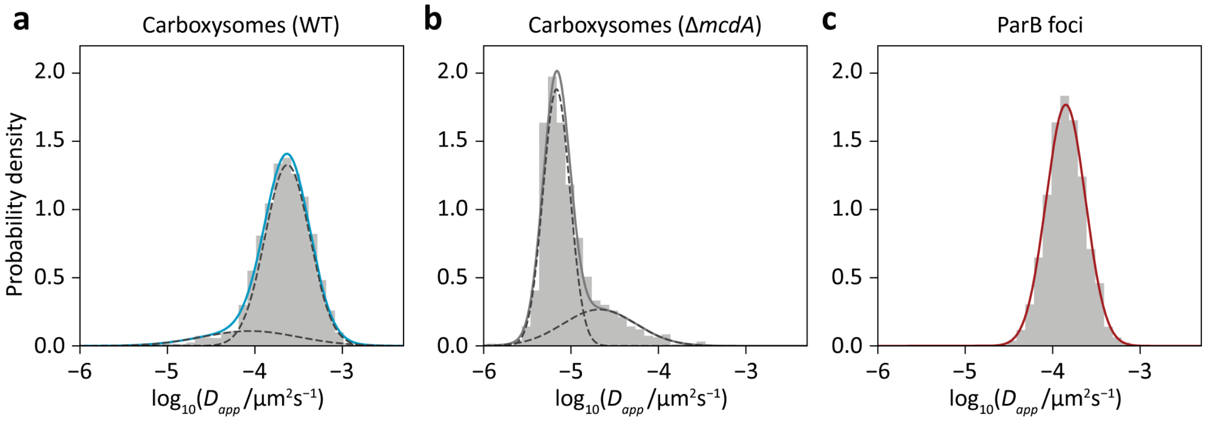


**d**

| | $D_{app}$ ($\mu m^2 s^{-1}$) | Fraction | $D_{app}$ ($\mu m^2 s^{-1}$) | Fraction |
|---|---|---|---|---|
| Carboxysomes (WT) | $(1.04 \pm 0.14) \times 10^{-4}$ | 17 ± 5% | $(2.40 \pm 0.05) \times 10^{-4}$ | 83 ± 5% |
| Carboxysomes (Δ*mcdA*) | $(0.07 \pm 0.01) \times 10^{-4}$ | 73 ± 2% | $(0.23 \pm 0.02) \times 10^{-4}$ | 27 ± 2% |
| ParB foci | $(1.4 \pm 0.02) \times 10^{-4}$ | 100% | | |

**Supplementary Fig. 2**: **Analysis of carboxysome and ParB focus diffusion**. Probability density distributions of apparent diffusion coefficients, $D_{app}$, for carboxysome foci in **a** WT (*N* = 3630 trajectories) and **b** Δ*mcdA* cells (*N* = 1896 trajectories), and **c** for ParB-mNG foci (*N* = 1284 trajectories) based on tracks collected from 1-s time-lapse imaging. In each plot, the $D_{app}$ distribution is fit by a one- or two-state Gaussian Mixture Model (GMM) (solid line). Dashed lines indicate the fits of the individual states. **d.** Values for average $D_{app}$ and population fraction (mean ± SD from 100 bootstrapped samples).

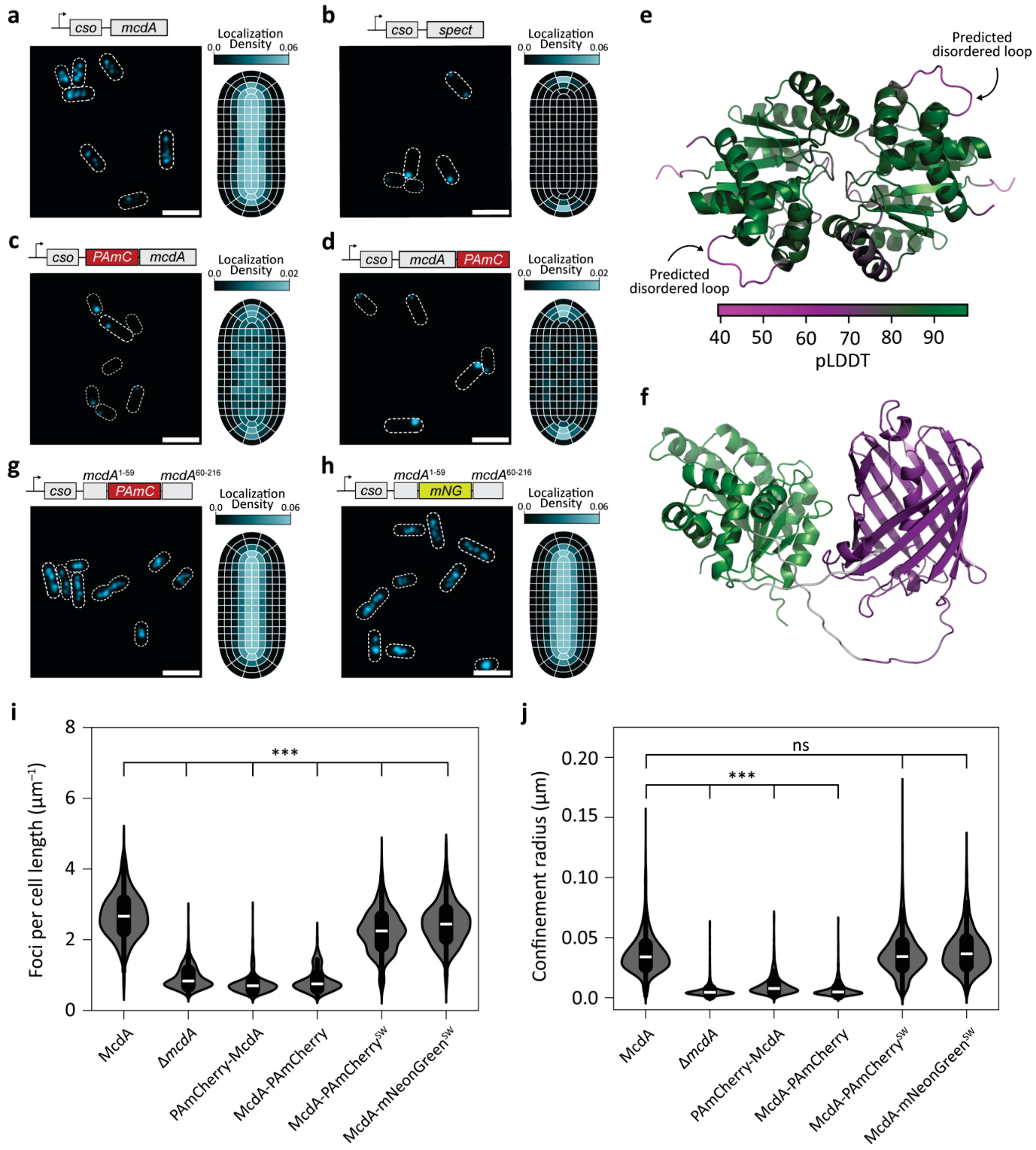


**Supplementary Fig. 3: Analysis of functional McdA fluorescent fusions**. **a-d**. Representative images of carboxysome localizations in **a** WT, **b** Δ*mcdA*, **c** PAmCherry-McdA, and **d** McdA-PAmCherry cells. The carboxysome signal (cyan) is from CbbS-mTQ. Dashed white outlines demarcate the cell outlines. Scale bars: 2 µm. Gene constructs (top) within the carboxysome (*cso*) operon. (Right) Carboxysome localization density heatmaps for each strain. **a** $N$ = 1061 cells, **b** $N$ = 1106 cells, **c** $N$ = 632 cells, **d** $N$ = 494 cells. **e.** McdA homodimer structure generated with AlphaFold2 (AF2). Color scale indicates the predicted Local Distance Difference Test (pLDDT). **f.** AlphaFold2-generated structure of McdA monomer (green) fused to PAmCherry (violet) in the sandwich orientation. **g-h.** Representative carboxysome localization for cells expressing **g** McdA-PAmCherry$^{SW}$ and **h** McdA-mNeonGreen$^{SW}$ fusions. Content as in **a-d**. **g** $N$ = 987 cells, **h** $N$ = 1052 cells. Scale bars: 2 µm. **i.** Number of carboxysome foci per unit of cell length for each strain. McdA $N$ = 1061 cells, $\Delta$*mcdA* $N$ = 1106 cells, PAmCherry-McdA $N$ = 871 cells, McdA-PAmCherry $N$ = 683 cells, McdA-PAmCherry$^{SW}$ $N$ = 1623 cells, McdA-mNeonGreen$^{SW}$ $N$ = 1052 cells. **j.** Confinement radius for carboxysome trajectories in each strain. McdA: $N$ = 3680 foci, $\Delta$*mcdA*: $N$ = 1896 foci, PAmCherry-McdA: $N$ = 950 foci, McdA-PAmCherry: $N$ = 1217 foci, McdA-PAmCherry$^{SW}$: $N$ = 2355 foci, McdA-mNeonGreen$^{SW}$: $N$ = 2781 foci. In **i** and **j**, the white lines are the medians, and the black cores are the interquartile range. *** denotes $p < 0.001$ by Kruskal-Wallis' test. ns denotes $p > 0.05$: McdA-PAmCherry$^{SW}$ $p = 0.54$, McdA-mNeonGreen$^{SW}$ $p = 0.23$. Statistical tests were done on the mean of means from 10 bootstraps, each randomly sampling 100 data points.

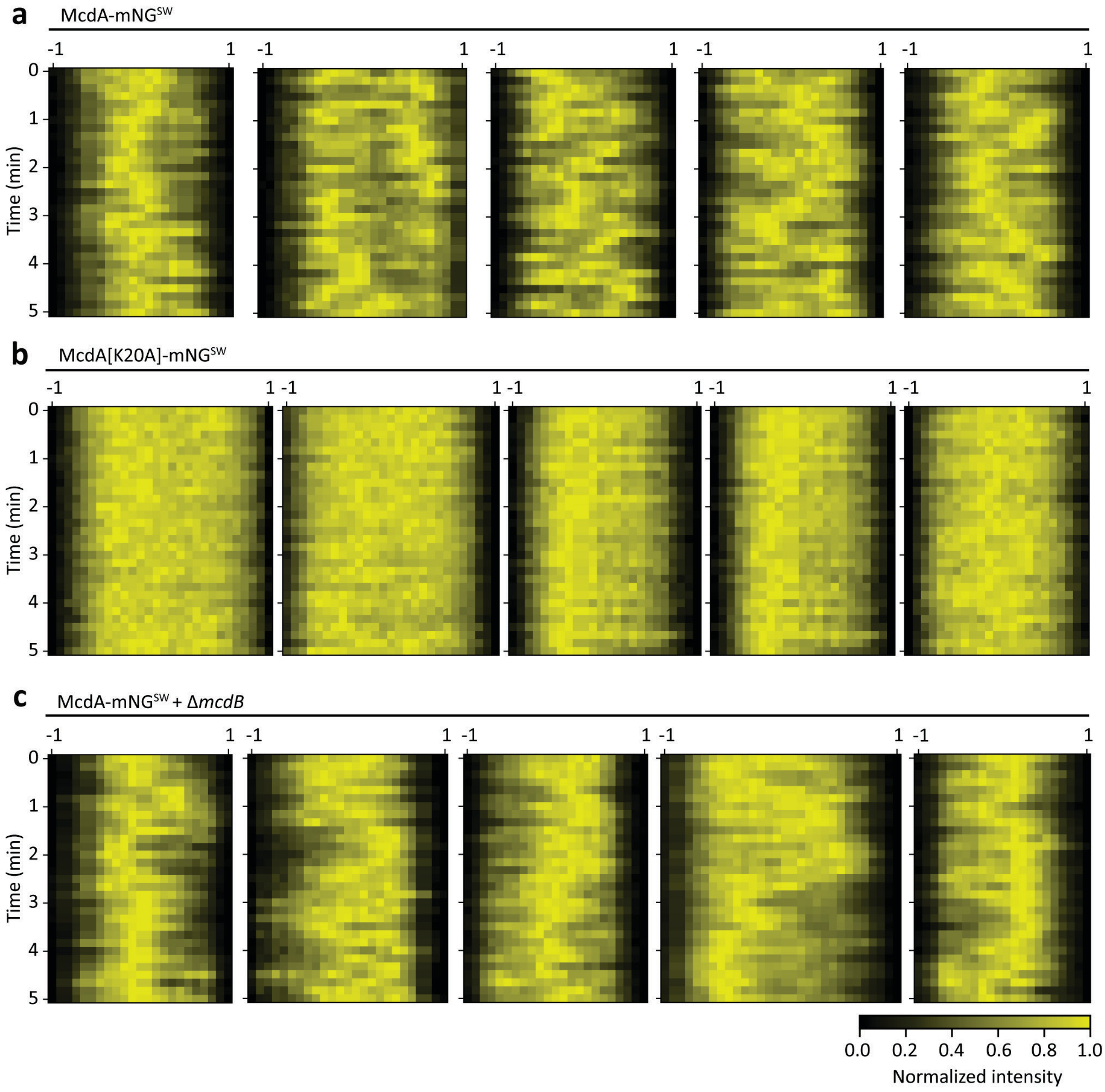


**Supplementary Fig. 4: Examination of bulk McdA localization**. Representative single-cell kymographs for **a** McdA-mNG$^{SW}$ in WT cells, **b** McdA[K20A]-mNG$^{SW}$ in WT cells, and **c** McdA-mNG$^{SW}$ in Δ*mcdB* cells. Time-lapse frames were collected every 10 seconds for 5 minutes. Profiles are normalized to the maximum and minimum intensities within the corresponding time point.

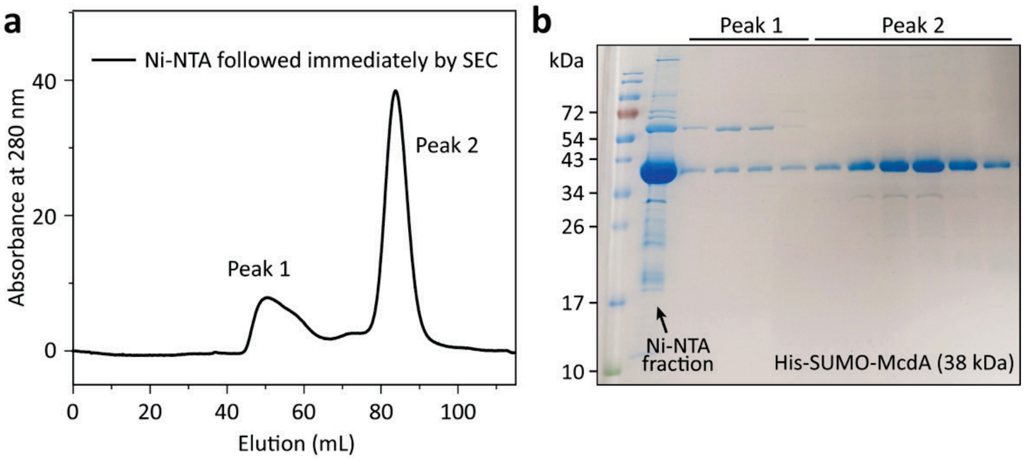


**Supplementary Fig. 5: Purification of *H. neapolitanus* His-SUMO-McdA**. **a.** Size-exclusion chromatogram of His-SUMO-McdA. Peaks indicate the eluting populations. **b.** SDS-PAGE of elution fractions collected from (**a**). Peak 2 is a pure band at ~38 kDa, consistent with His-SUMO-McdA.

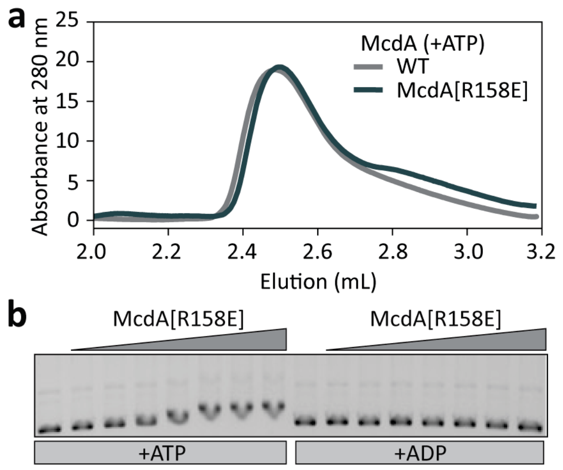


**Supplementary Fig. 6**: **Biochemical analysis of McdA[R158E]. a.** Gel-filtration analysis of purified His-SUMO-McdA and His-SUMO-McdA[R158E] in the presence of ATP. **b.** Electrophoretic mobility shift assay (EMSA) with increasing concentrations (0, 0.5, 1, 2.5, 5, 10, 12.5, 14 µM) of His-SUMO-McdA[R158E]. 1 mM of ATP or ADP was used.

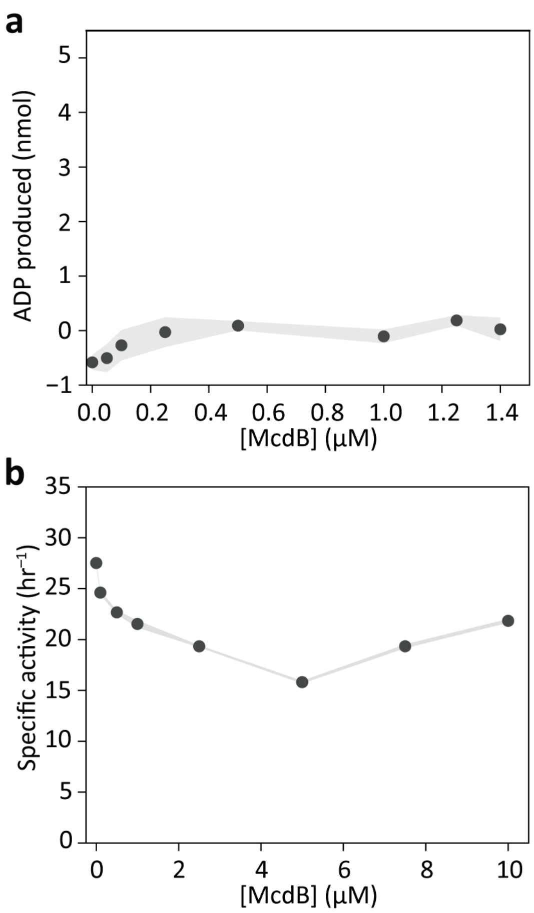


**Supplementary Fig. 7: ATPase activity of McdA in the presence of excess McdB. a.** ATPase activity of McdB with DNA (0.1 mg/mL). [ATP] was kept fixed at 1 mM. Amount (nmol) of ADP produced in reaction after 2 hr. Shading indicates the SEM of three independent experiments. **b.** Specific activity (turnover rate: nmol ADP produced/hr vs. nmol McdA) of His-SUMO-McdA (concentration fixed at 0.25 µM) measured as a function of McdB concentration (0 – 10 µM). Shading indicates the error on the linear regression.

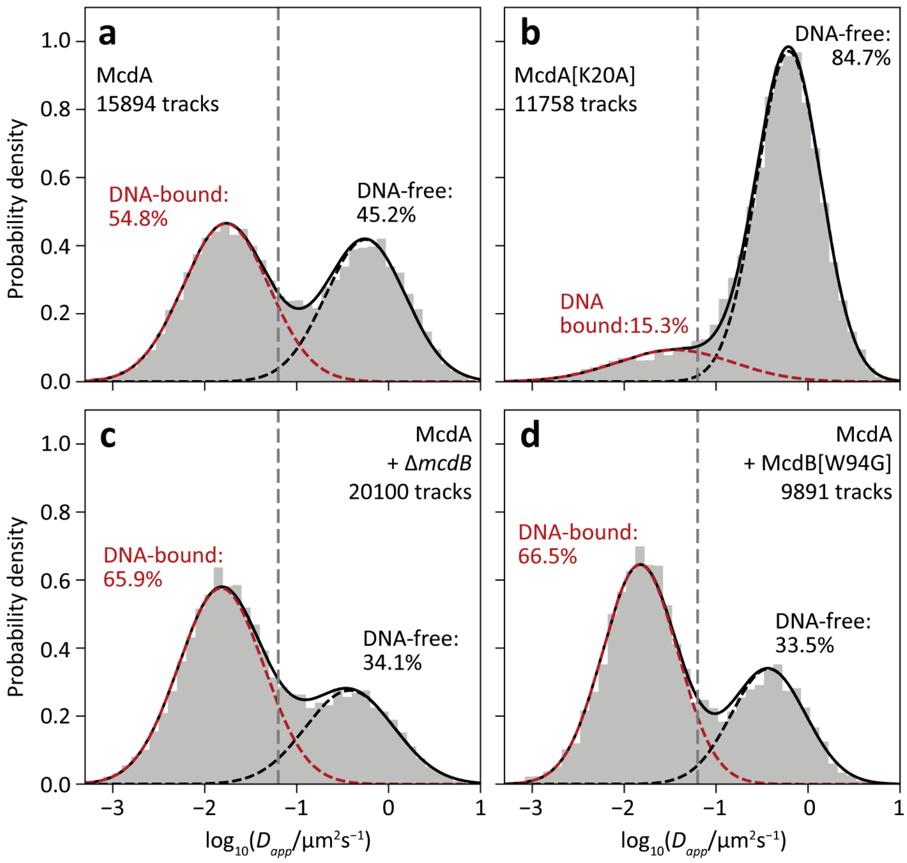


**Supplementary Fig. 8: Diffusion analysis of single-molecule trajectories of McdA and its mutants collected with 20-ms camera integration time**. Probability density distributions of apparent diffusion coefficients, $D_{app}$, for **a** McdA-PAmCherry$^{SW}$ in WT cells, **b** McdA[K20A]-PAmCherry$^{SW}$ in WT cells, **c** McdA-PAmCherry$^{SW}$ in Δ*mcdB* cells, and **d** McdA-PAmCherry$^{SW}$ in McdB[W94G] cells based on tracks collected with 20-ms camera integration times for all strains imaged in this study. Also shown in each plot is the fit to a two-state Gaussian Mixture Model (GMM) (solid black line). Colored dashed lines indicate the individual states of the GMM fit, colored-coded by the assigned biological state. Only tracks ≥ 5 frames are included. Data are from three or more biological replicates for each strain. The dashed gray line is the theoretical lower bound for measurable $D_{app}$ based on localization uncertainty from a fixed sample (Supplementary Fig. 9).

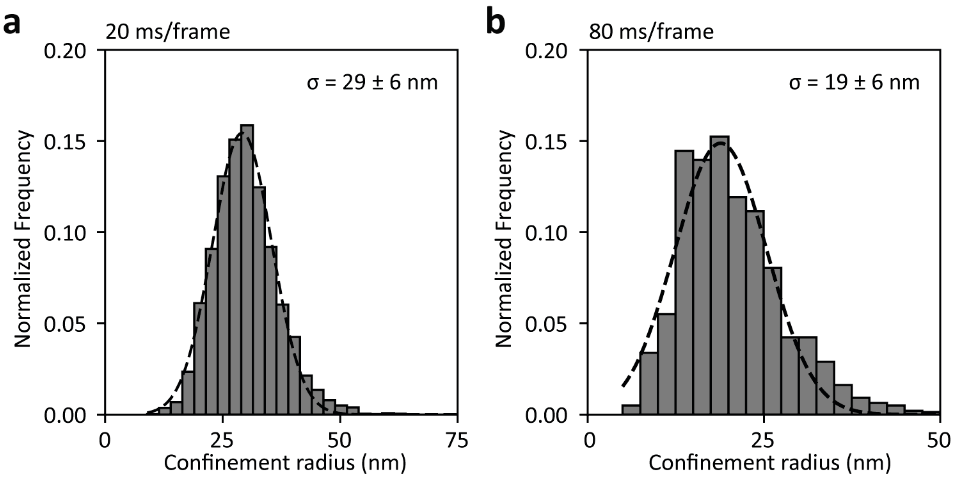


**Supplementary Fig. 9: Localization uncertainty associated with consecutive detections of single PAmCherry molecules in fixed *H. neapolitanus* cells**. Single molecules were imaged with **a** 20-ms and **b** 80-ms camera integration times. The confinement radii of $N$ = 3827 tracks with ≥ 5 localizations were calculated by averaging the distance of every localization to the centroid (average position) of the track. Also shown is the Gaussian model fit (dashed line) of the confinement radius histogram, $p(r)$, using Equation 4.

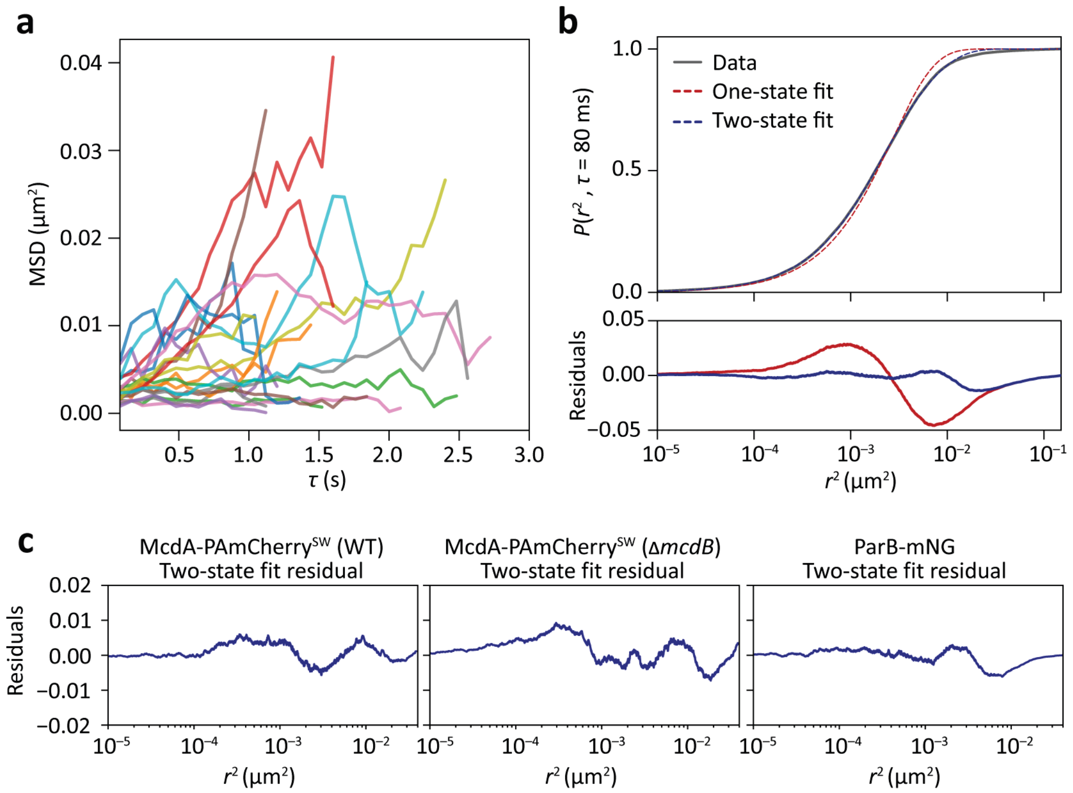


**Supplementary Fig. 10: Nucleoid-bound McdA-PAmCherry$^{SW}$ dynamics are heterogeneous.** **a.** MSD versus time-lapse, $\tau$, curves for 20 randomly sampled McdA-PAmCherry$^{SW}$ trajectories from Δ*mcdB* cells imaged continuously at 80-ms integration times. Curves show immobile, mobile, and switching behavior. **b.** Top: Cumulative distribution function (CDF) of squared displacements with $\tau$ = 80 ms for continuous McdA-PAmCherry$^{SW}$ trajectories in Δ*mcdB* cells (*N* = 4487 tracks across three datasets). One-state (red dashed line) and two-state (blue dashed line) models were fit to the CDF (see Methods). Bottom: Residuals for the one-state (red line) and two-state (blue line) fits. **c.** Residuals for the two-state fits to the CDFs in Figure 5c for McdA-PAmCherry$^{SW}$ trajectories collected from WT (left) and Δ*mcdB* (center) cells and ParB-mNG foci (right) at 80-ms integrations and 140-ms dark-time delays.

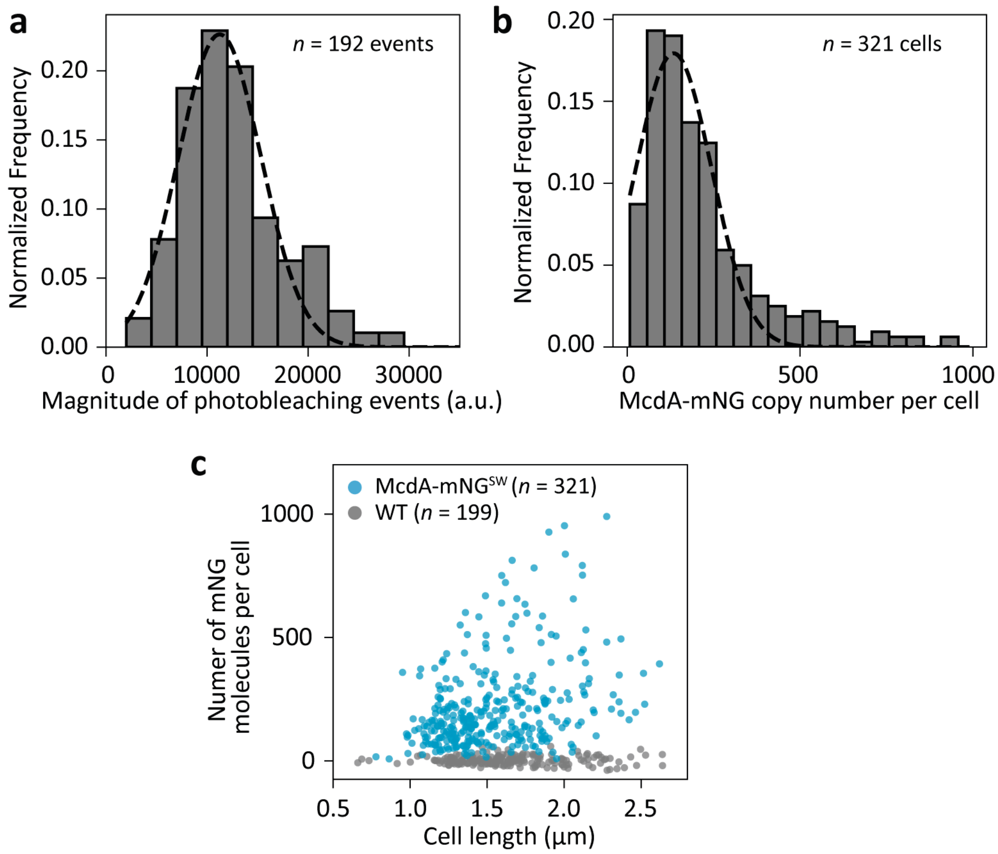


**Supplementary Fig. 11: Fluorescence-based quantification of McdA-mNG$^{SW}$ copy number in *H. neapolitanus* cells. a.** Distribution of the magnitude of fluorescence intensity loss following each single-step photobleaching of mNG-FliN. *N* = 192 photobleaching events. The mean of the Gaussian fit (dashed line) gives the fluorescence intensity of a single mNG molecule in *H. neapolitanus*: 11,245 ± 4094 counts. **b.** Distribution of the number of McdA-mNG$^{SW}$ copies per *H. neapolitanus* cell. *n* = 321 cells. The mean of the Gaussian fit (dashed line) gives the average McdA copy number per cell: 195 ± 79 McdA-mNG$^{SW}$ molecules. **c.** Calculated number of McdA-mNG$^{SW}$ molecules per cell as a function of cell length for control (wildtype cells expressing no mNG; *n* = 199) and *mcdA-mNG$^{SW}$* cells (*n* = 321).

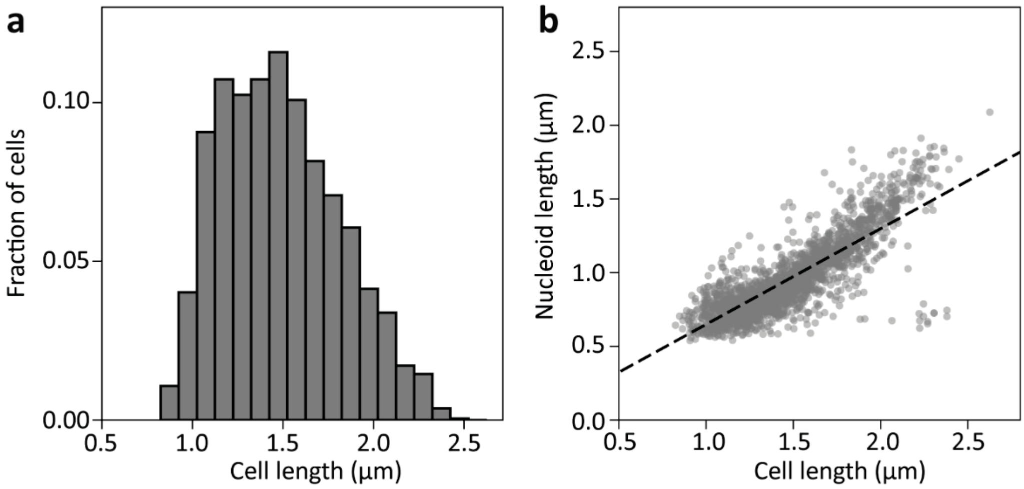


**Supplementary Fig. 12: *H. neapolitanus* cell and nucleoid lengths**. **a.** Distribution of WT cell lengths ($N$ = 1864 cells). The average cell length is 1.49 ± 0.33 μm. **b.** Relationship between cell length, $L_{cell}$, and nucleoid length, $L_{nucleoid}$ ($N$ = 1864 cells). Each point is an individual cell. Dashed line indicates a linear fit to the data: $L_{nucleoid} = 0.65L_{cell}$.
